# Essential regulation of heparan sulfate proteoglycan signalling controls cell behaviour to support cardiac development

**DOI:** 10.1101/2025.07.23.666364

**Authors:** Andia N. Redpath, Irina-Elena Lupu, Louis Haffreingue, Quang Dang, Ian R. McCracken, Tamara Carsana, Toin H. van Kuppevelt, Joaquim Miguel Vieira, Nicola Smart

**Affiliations:** Institute of Developmental and Regenerative Medicine, Department of Physiology, Anatomy & Genetics, University of Oxford, Oxford OX3 7TY, UK; 2High-Tech Center, Vinmec International Healthcare System, Hanoi, Vietnam; Department of Biochemistry, Radboud University Medical Center, Nijmegen, The Netherlands; School of Cardiovascular & Metabolic Medicine & Sciences, King’s College London, London SE5 9NU, UK

## Abstract

Heparan sulfate proteoglycan (HSPG)-dependent signalling is essential for cardiac development, yet how HSPGs are regulated in the forming heart is unknown. Here, we show that the extracellular heparan sulfate modifier, SULF1, expressed by the epicardium, dynamically regulates HSPG sulfation to fine-tune the magnitude and duration of signalling to impact cell fate and cardiac morphogenesis. Single cell genomics and lineage tracing studies reveal *Sulf1* to be strongly co-expressed with *Wt1* in the epicardium, with reduction of both coinciding with epithelial-mesenchymal transition and quiescence. CUT&RUN-seq revealed transcriptional control of *Sulf1* by WT1 which directly impacts essential HSPG-dependent downstream signalling. Our study highlights, for the first time, essential fine-tuning of HSPG-dependent signalling to modulate key processes for heart formation.

## Main

The outermost layer of the heart consists of specialised mesothelium, referred to as the epicardium, which originates from a transient structure called the proepicardium (PE) ^1, 2^. The epicardium fundamentally controls cardiogenesis, by producing a plethora of angio- and cardio-genic paracrine signals and extracellular matrix (ECM) components, and by contributing essential cell types ^3^. Key to myocardial invasion and contribution is the extensive proliferation of epicardial cells, followed by a highly regulated transformation, epithelial-mesenchymal transition (EMT), to give rise to epicardium-derived cells (EPDCs). Although the epicardium is quiescent in the adult, cardiac injury induces a degree of epicardial reactivation, which is necessary for full regeneration in lower vertebrates such as zebrafish ^4^. While partial epicardial reactivation in the mammalian heart augments cardiac repair, healing is limited by inadequate epicardial cell invasion ^5^. Delineating the mechanisms that control proliferation and EMT during development may reveal targets for enhancing epicardium-based repair.

Although EMT is reported to fall under the control of the Wilms’ Tumour 1 (WT1) transcription factor ^6^, touted as a ‘master regulator’ of epicardial biology, few direct targets of WT1 have been identified ^7^, and little is known of how EMT is controlled at the transcriptional level. During development, epicardial processes are efficiently stimulated by coordinated growth factor signalling. First, fibroblast growth factors (FGF) and Wnt family members (WNT) control initial formation ^8–12^, insulin-like growth factors (IGF) promote proliferation ^13^, while numerous factors induce EMT, including transforming growth factor β (TGF-β) ^14–16^, FGF ^16–18^, and platelet-derived growth factor (PDGF) ^16, 19^. A common feature of these pathways is their regulation by heparan sulfate proteoglycans (HSPG), that tightly control growth factor/morphogen signalling pathways in embryonic development ^20^. In brief, HSPGs may modulate signalling by binding growth factors to: provide an extracellular depot; recruit them to the cell surface in proximity to their corresponding receptors; or facilitate the formation of gradients essential for specification and migration. Membrane-bound HSPGs can act as co-receptors for numerous growth factors, including FGF ^21–25^ and TGF-β ^26–28^, by lowering receptor activation thresholds or affecting the duration of signalling ^29^. In heart development, several loss-of-function studies revealed roles for HSPGs in cardiac lineage specification ^30, 31^, outflow tract and atrioventricular valve formation ^32, 33^, and ventricular septation ^34^. These studies evidenced a critical requirement for HSPG proteins in cardiac specification and morphogenesis, albeit only a minor fraction of vertebrate HSPGs were explored.

HSPGs comprise a core protein, Xyl-Gal-Gal linker, with IdoA/GlcA-GlcN or -GlcNAc saccharide repeats that are sulfated in various positions. Diversity in the extent and position of HS chain sulfation determines capacity for interaction with growth factors and receptors ^20^, meaning that sulfation status adds another layer of complexity to signalling pathway regulation. The 6-*O*-sulfate moiety, key to promoting ternary complex formation of, for example, FGF-FGFR-HS ^35^, is added to C6 by heparan sulfate 6-*O*-sulfotransferases. Deficiency of HS-generating enzymes causes a range of cardiac abnormalities during development, in some cases explained by a disruption in HS-dependent FGF or BMP signalling ^36, 37^. Countering the actions of 6-*O*-sulfotransferases, 6-*O*-endosulfatases, SULF1 and SULF2, remove 6-*O* sulfate moieties from HS chains, and are central to the development of the dynamic ‘sugar code’ post-synthetically, and to temporal and regional sulfation patterning in organs ^38, 39^. Knock-out of individual 6-*O*-endosulfatases has not consistently manifested developmental abnormalities ^40–45^, due to non-penetrance and functional redundancy. However, genetic disruption of either isoform impacted neovascularisation via HS-binding VEGF-A_164_ and heart repair following myocardial infarction ^46^, leading to increased scarring and cardiac dysfunction. The detection of *Sulf1* in the injury re-activated epicardium ^47^ establishes the potential for HSPG signalling to profoundly control epicardial cell behaviour post-myocardial infarction, and to influence its contribution to regeneration. Given that extracellular endosulfatases may determine epicardial responses to incoming signals and consequently influence epicardial behaviour over the course of heart development, and possibly in disease, the regulation of 6-*O*-endosulfatases and their actions warrants investigation.

In the present study, we systematically profiled 6-*O*-endosulfatases and target HSPGs in the embryonic mouse heart. We identified cell type-specific expression patterns for *Sulf1* and *Sulf2*, that changed over the course of development, unrelated to lineage or embryonic origin. We reveal *Sulf1* as the major 6-*O*-endosulfatase expressed in the epicardium, and accordingly, defects in epicardial formation in conditional *Sulf1* knock-out mice were detected, that partly phenocopied *Wt1* knockout embryos. Indeed, we show that *Sulf1* expression closely correlates with *Wt1*, with both downregulating upon EMT and quiescence, and elucidated that WT1 regulates *Sulf1* levels in the epicardium, via binding to a *Sulf1* intronic enhancer. Finally, we quantitatively inferred HSPG-dependent signalling during EMT and found that key factors promoting epicardial proliferation and EMT, FGF2 and TGF-β1, were significantly impacted by the WT1-*Sulf1* regulatory interaction. Collectively, this work presents a mechanism linking transcriptional control of HS sulfation to modulation of growth factor signalling and instruction of cellular responses, to correctly orchestrate heart development. Such insights may be important for therapeutically enhancing epicardial cell proliferation and EMT in the adult heart for regeneration.

## Results

### HSPGs and their modifiers demonstrate distinct expression in the developing heart

Despite their decisive influence over the signalling that underlies cardiac morphogenesis, the expression of core HSPGs and the enzymes that give rise to distinct spatiotemporal sulfation patterning have not been well documented in the developing heart. We, therefore, systematically profiled HSPG expression at three key embryonic stages E10.5, E13.5 and E17.5. We analysed the published E10.5 mouse heart scRNA-seq dataset (^48^; Extended Data Fig. 1A), and generated E13.5 (^49^; Extended Data Fig. 1B) and E17.5 (Extended Data Fig. 1C, D) heart scRNA-seq datasets with epicardial lineage tracing (Extended Data Fig. 1E). HSPGs are expressed in mammalian cells and may be divided into ‘full-time’ and ‘part-time’ categories, with at least 95% of the heparan sulfate accounted for by ‘full-time’ HSPGs ^50^. ‘Full-time’ HSPGs include the membrane-bound syndecans (*Sdc1*-*4*) and glypicans (*Gpc1*-*Gpc6*), ECM/basement membrane localised perlecan (*Hspg2*), agrin (*Agrn*) and collagen XVII (*Col18a1*), and secretory vesicle localised serglycin (*Srgn*). ‘Part-time’ HSPGs include membrane-bound betaglycan (*Tgfbr3*), epican (*Cd44*), and neuropilin-1 (*Nrp1*), and ECM localised testicans (*Spock1-3*) ^29, 51^. The 20 HSPGs we interrogated not only demonstrated heterogeneous expression profiles between cardiac cell types, but also between embryonic stages (Extended Data Fig. 1F), with special exception of *Srgn* largely expressed in the resident leukocyte population, and *Nrp1* ubiquitously expressed in the heart (excluding lymphatic endothelial cells (LECs); Extended Data Fig. 1F). *Col18a1* was primarily expressed in non-myocyte populations. *Gpc2*, *Gpc5, and Spock1-3* were not detected, in keeping with failure to detect these HSPG types in embryonic and adult mouse hearts ^52, 53^. We noted *Sdc2*, *Sdc4*, *Nrp1*, *Gpc3* and *Col18a1* consistently expressed in epicardial cells, while *Sdc4* was downregulated and *Gpc6* upregulated in epicardium-derived cells (Extended Data Fig. 1F). These results indicate that HSPGs are dynamically expressed, with cell-type specific expression, and changing tissue composition contributing to HSPG diversity in the developing embryonic heart.

We next explored expression of 6-*O*-sulfation enzymes throughout the heart. Despite some variation in expression over the three embryonic stages, the sulfotransferase isoforms *Hs6st1* and *Hs6st2* were mainly expressed by epicardial cells and cardiomyocytes (Extended Data Fig. 2A). *Hs6st3* expression was not detected in these datasets (Extended Data Fig. 2A), supported by the general absence of this isoform in embryonic and adult hearts ^53, 54^. Finally, we determined cardiac expression of 6-*O*-endosulfatases, which refine organ-specific sulfation patterning through selective 6-*O* desulfation ^38, 39^. We detected enriched *Sulf1* expression in the epicardial clusters at all stages examined, with sporadic and low expression also in endothelial populations, and an increased expression within mesenchyme over the course of development (Fig. 1A-C). In contrast, *Sulf2* was broadly expressed throughout development, with variable levels in most cardiac cell types (Fig. 1A-C). The dynamic profile of *Sulf1*, increasing from E11.5 to E15.5 before declining, was quantitatively confirmed in whole embryonic heart lysates, and is in contrast to *Sulf2* expression, which was unchanged (Extended Data Fig. 2B).

**Fig. 1.**
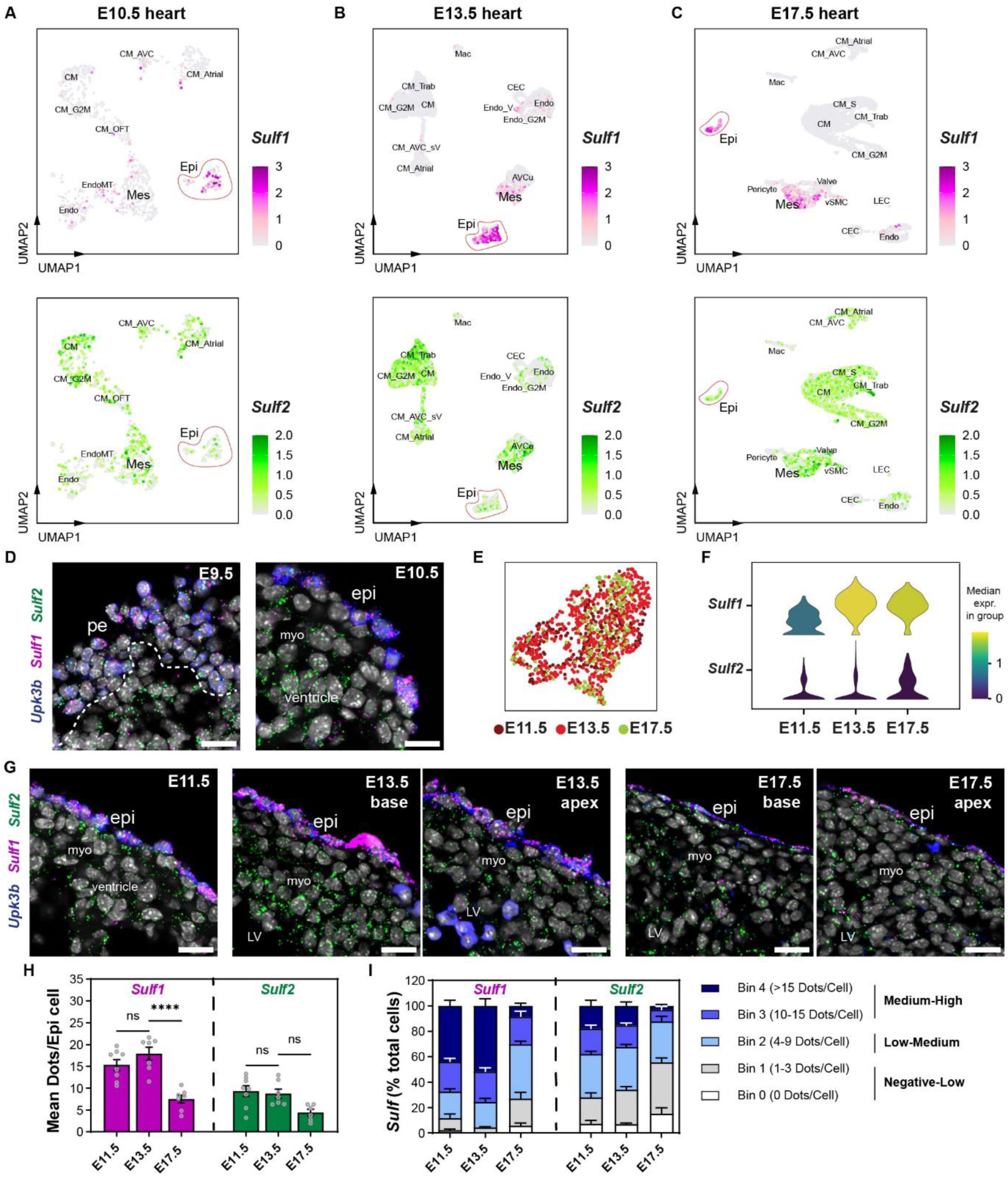
*Sulf1* and *Sulf2* demonstrate distinct expression profiles in the developing heart. (A) Uniform manifold approximation and projection (UMAP) feature plot showing expression of *Sulf1* and *Sulf2*, in individual cells of E10.5 hearts (total 928 cells, n = 2 batches). (B) UMAP feature plot showing expression of *Sulf1* and *Sulf2*, in individual cells of E13.5 hearts (total of 6,939 cells, n = 3 hearts). (C) UMAP feature plot showing expression of *Sulf1* and *Sulf2*, in individual cells of E17.5 hearts (total of 6,341 cells, n = 1 heart). (D) Fluorescence ISH on cryosections of E9.5 and E10.5 embryos for *Upk3b*, *Sulf1* and *Sulf2*, showing expression of *Sulf1* in the PE and newly-formed epicardium. Scale bars: 20 µm. Images representative of n = 4 embryos for E9.5 and n = 7 embryos for E10.5. (E) UMAP plot showing integrated epicardial clusters (1093 cells), combining three datasets from E11.5, E13.5 and E17.5 hearts. (F) Stacked violin plot showing the expression of *Sulf1* and *Sulf2* in the epicardial clusters across developmental stages. High median expression is indicated in yellow. (G) Fluorescence ISH on cryosections of E11.5, E13.5 and E17.5 hearts for *Upk3b*, *Sulf1* and *Sulf2*, showing expression of *Sulf1* in the epicardium, while widespread expression of *Sulf2* is detected in the myocardium. Scale bars: 20 µm. Images representative of n = 9 hearts/embryos for E11.5, n = 8 hearts for E13.5, and n = 8 hearts for E17.5. (H) Bar plots showing quantification of *Sulf1* and *Sulf2* expression in ventricular epicardial cells (of Fluorescence ISH images). Data are represented as mean ± SEM (n = 6-8 hearts). One-Way ANOVA and Šídák multiple comparison test (****) P <0.0001 and ns, not significant. (I) Percentage of epicardial cells expressing *Sulf1* and *Sulf2*, grouped into 5 bins based on the number of dots (individual RNA molecules) per cell. Bin 0, 0 dots/cell; Bin 1, 1-3 dots/cell; Bin 2, 4-9 dots/cell; Bin 3, 10-15 dots/cell; Bin 4, >15 dots/cell. Data are represented as mean ± SEM (n = 6-8 hearts). *AVCu, atrioventricular cushion; CEC, coronary endothelial cells; CM, ventricular cardiomyocytes; CM_Atrial, atrial cardiomyocytes; CM_AVC, atrioventricular canal cardiomyocytes; CM_AVC_sV, atrioventricular canal and sinus venosus cardiomyocytes; CM_OFT, outflow tract cardiomyocytes; CM_S, S-G2M phase progressing cardiomyocytes; CM_Trab, trabecular cardiomyocytes; Endo, endocardial cells; Endo_V, valvular endothelial and endocardial cells; EndoMT, endocardial-to-mesenchymal transition; Epi, epicardium; G2M, proliferating; LEC, lymphatic endothelial cells; Mac, macrophages; Mes, mesenchyme; myo, myocardium; pe, proepicardium; Valve, valve mesenchyme; vSMC, vascular smooth muscle cells*.

Consistent with the epicardial enrichment, we detected *Sulf1* expression as early as E9.5 in the proepicardium (Fig. 1D), the origin of epicardial progenitors, and next in the newly formed epicardial layer (Fig. 1D, Extended Data Fig. 2C; E10.5). To enable comparison once the epicardium is established, we utilised an E11.5 scRNA-seq dataset ^55^, and following data integration (Fig. 1E), we elucidated a temporal pattern in *Sulf1* expression levels (Fig. 1F). Indeed, peak epicardial expression of *Sulf1* was noted at E13.5, and diminished by E17.5 (Fig. 1G-I). Although notably at E17.5, expression persists at a higher level in the atrial epicardium, in comparison to that of the ventricles (Extended Data Fig. 2D-F). Meanwhile, a smaller subset of epicardial cells co-expressed *Sulf2*, with evidently fewer transcripts than *Sulf1* (Fig. 1G-I) and, in accordance with the scRNA-seq, we observed cardiomyocytes as the major source of *Sulf2* in the developing heart (Extended Data Fig. 2C, C’). Altogether, these findings demonstrate that 6-*O*-endosulfatase isoforms have distinct expression in the embryonic heart, revealing restricted overlap in some cells. We identified an abundant and localised expression of *Sulf1* in the epicardium, suggesting it may perform extracellular HS modifications of potential targets syndecan-4, neuropilin-1 and glypican-3 for epicardial signalling regulation.

### *Sulf1* and *Sulf2* are expressed in a subset of non-myocyte cells irrespective of origin

Given the accumulation of *Sulf1/2*-expressing mesenchyme over the course of development (Fig. 1A-C), we sought to determine whether endosulfatase expression in cardiac mesenchyme, fibroblasts and mural cells was specific to their epicardial origin. We utilised the epicardial-lineage tracing capabilities of *Wt1^CreERT2/+^;Rosa26^tdTomato^*mice (Extended Data Fig. 1E), with Tamoxifen administered before embryonic stage E11.5, to label the newly formed epicardium (Fig. 2A) and minimise labelling of sinus venosus-derived *Wt1*-expressing endothelium ^2, 56^. The tdTomato-labelled epicardial layer expressed higher *Sulf1* compared to its early derivatives marked by *tdTomato* and *Postn* co*-*expression (Fig. 2B). *Sulf*-expressing epicardium-derived mesenchymal cells were localised in high EMT activity regions such as the atrioventricular groove and apical subepicardium (Fig. 2B, Extended Data Fig. 3A). At E17.5, we observed an upregulation of *Sulf1* in the tdTomato-labelled mesenchymal cells (*Dpt* positive; Fig. 2C) that had invaded furthest into the myocardium (Extended Data Fig. 3B, C). Finally, *Sulf1* and *Sulf2* were detectable, albeit at strikingly low levels, in epicardium-derived vascular smooth muscle cells (vSMCs; *Rgs5* and *Myh11* positive; Fig. 2D, Extended Data Fig. 3D). To assess the origin of mesenchyme and mural cells that express 6-*O*-endosulfatases, we analysed scRNA-seq (Fig. 2E, Extended Data Fig. 3E) and flow cytometry-based RNA *in situ* hybridization (PrimeFlow RNA assay; Fig. 2F-I) of E17.5 hearts. Mesenchymal and mural cells were assigned as epicardium-derived (tdTomato positive) or originating from an alternative source (tdTomato negative), likely the endocardium ^57, 58^. 40-60% of mesenchymal cells and vSMCs expressed 6-*O*-endosulfatases irrespective of their origin (Fig. 2E-I). However, notably, *Sulf1* was more abundantly expressed (transcripts per cell) in epicardium-derived mesenchymal cells compared with mesenchyme originating from other sources (Fig. 2F, G).

**Fig. 2.**
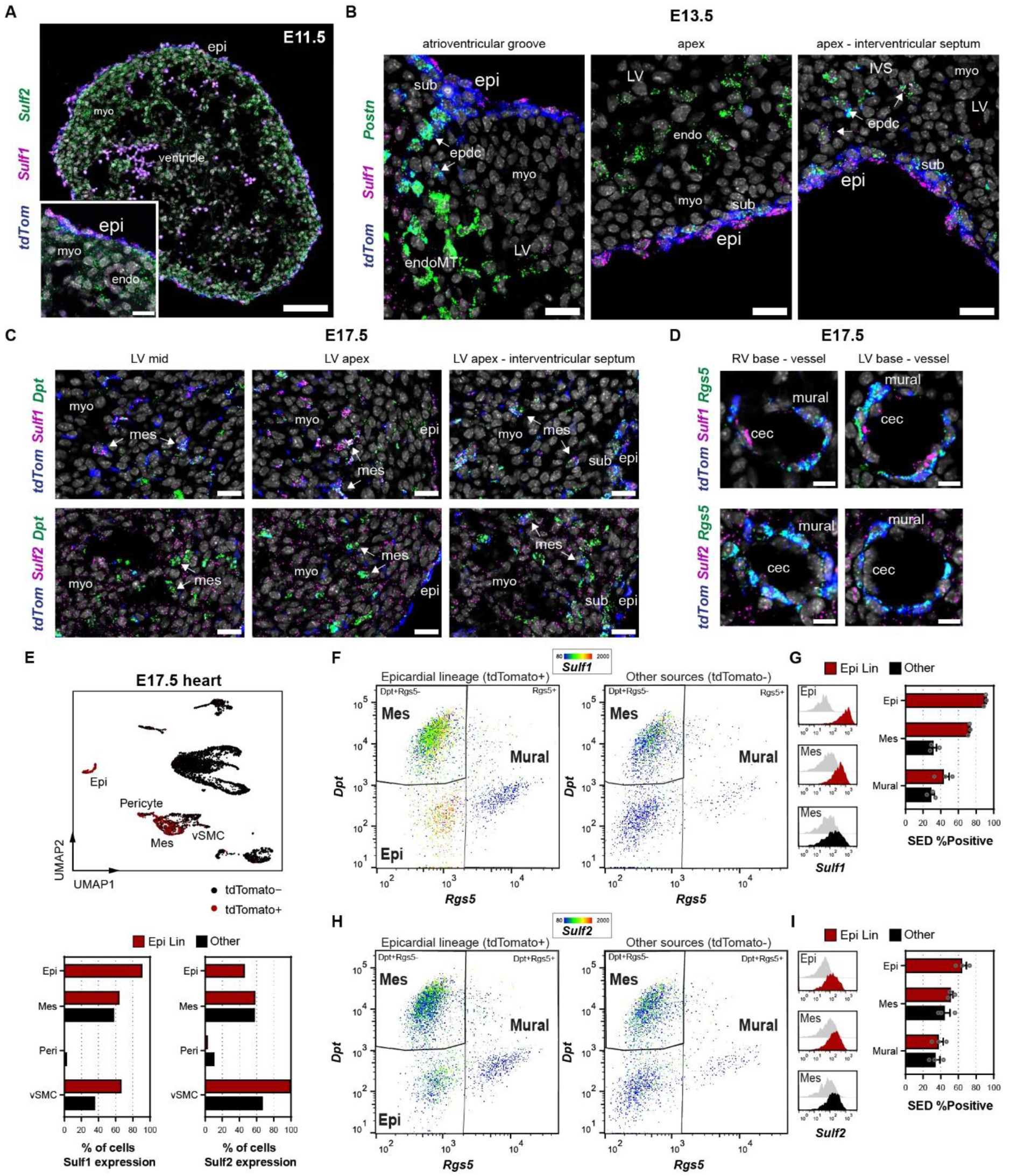
*Sulf1* and *Sulf2* are expressed in a subset of cardiac mesenchyme and perivascular cells irrespective of origin. (A) Fluorescence ISH of E11.5 heart cryosections for *tdTomato*, *Sulf1*, and *Sulf2*, showing expression of *Sulf1* enriched in the tdTomato-labelled epicardium of *Wt1*^CreERT2^;Rosa26*^tdTomato^* embryos induced at E9.5. Scale bars: 100 µm and inset 20 μm. Images representative of n = 6 embryos/hearts. (B) Fluorescence ISH of E13.5 heart cryosections for *tdTomato*, *Sulf1* and *Postn*, showing expression of *Sulf1* in epicardial-lineage subepicardial mesenchymal cells. Scale bars: 25 µm. Images representative of n = 3 hearts. (C) Fluorescence ISH of E17.5 heart cryosections for *tdTomato*, *Sulf1*, *Sulf2* and *Dpt*, showing expression of *Sulf1* and *Sulf2* in epicardium-derived mesenchymal cells invading the myocardium. Scale bars: 20 µm. Images representative of n = 5 hearts. (D) Fluorescence ISH of E17.5 heart cryosections for *tdTomato*, *Sulf1*, *Sulf2* and *Rgs5*, showing expression of *Sulf1* and *Sulf2* in epicardium-derived mural cells lining coronary vessels. Scale bars: 10 µm. Images representative of n = 3 hearts. (E) UMAP plot showing *tdTomato* positive cells in the E17.5 heart (total of 739 *tdTomato* positive cells). Populations expressing *Sulf1* and *Sulf2*, as a proportion of *tdTomato* positive epicardial lineage (Epi Lin) or *tdTomato* negative non-epicardial lineage (Other) mesenchymal and mural cells. (F-G) Flow cytometry analysis of E17.5 hearts, and corresponding quantification of percentage positive cells for *Sulf1*. The epicardial lineage selected by gating CD31^−^cTnT^−^tdTomato^+^ while non-epicardial selected by gating CD31^−^cTnT^−^tdTomato^−^ (other sources), with subsequent gating of mesenchymal cells (Dpt+Rgs5-) and mural cells (Rgs5+). (F) Colour mapping displays expression levels of *Sulf1*, showing highest expression in epicardial cells followed by epicardium-derived mesenchymal cells. (G) Fluorescence minus one (FMO) control shown (grey). Data are represented as mean ± SEM (n = 3 hearts). (H-I) Flow cytometry analysis of E17.5 hearts, and corresponding quantification of percentage positive cells for *Sulf2*. The epicardial lineage selected by gating CD31^−^cTnT^−^tdTomato^+^ while non-epicardial selected by gating CD31^−^cTnT^−^tdTomato^−^ (other sources), with subsequent gating of mesenchymal cells (Dpt+Rgs5-) and mural cells (Rgs5+). (I) Colour mapping displays expression levels of *Sulf2*, showing highest expression in epicardial and epicardium-derived mesenchymal cells. (G) Fluorescence minus one (FMO) control shown (grey). Data are represented as mean ± SEM (pooled 2 hearts per replicate; n = 3). *epi, epicardium; myo, myocardium; endo, endocardium; sub, subepicardium; epdc, epicardium-derived cells; endoMT, endocardial-to-mesenchymal transition; LV, left ventricle; IVS, interventricular septum; mes, mesenchymal cells; RV, right ventricle; cec, coronary endothelial cells*; *vsmc, vascular smooth muscle cells*.

*Sulf1* was previously implicated in the regulation of endothelial cells (ECs) during vascular development and ischaemic heart repair ^46, 59^. However, when profiling expression in the developing heart, we observed only modest expression in minor subsets within the coronary endothelial and endocardial clusters (Extended Data Fig. 3F-H). Specifically, we detected *Sulf1*, co-expressed with *Sulf2,* in coronary endothelial cells (CECs) with arterial macrovascular identity (vSMC coverage) located at the base of the heart (Extended Data Fig. 3D, H, I), and a subset of atrioventricular valve endothelial cells (VEC) directly facing blood flow (Extended Data Fig. 3H, J-K’). These atrioventricular VECs originate from the endocardium, demonstrated by their expression of *tdTomato* (Extended Data Fig. 3L), in temporally induced BmxCreERT2;Rosa26^tdTomato^-lineage labelled ECs ^60^. Potentially reflecting a distinct transcriptional driver, only *Sulf2* was detected in LECs (Extended Data Fig. 3H). These results demonstrate restricted expression pattern of 6-*O*-endosulfatase isoforms, in stark contrast to the pan-cardiac EC expression previously reported ^46, 59^.

### Dysregulated 6-*O*-endosulfatase expression impairs heart development

WT1 expression, which is largely reflective of epicardial activity during development, is most abundant in the newly formed epicardium at E11.5 ^2^, and its epicardial functions are essential for cardiac development ^6, 61, 62^. In contrast to *Sulf2*, epicardial *Wt1* and *Sulf1* expression decrease in levels as embryonic development progresses; *Wt1* downregulates after E11.5 ^2^, and *Sulf1* downregulates after E13.5 (Fig. 1D-H), to coincide with reduced epicardial proliferation (Extended Data Fig. 4A, B). We bred heterozygous carriers of the *Wt1-CreERT2* knock-in, an allele null for WT1 ^6^, to generate *Wt1* knockout (KO) embryos (Wt1^CreERT2/ CreERT2^; Fig. 3A). *Wt1* KO hearts demonstrated significant disruption, with few epicardial cells lining the E11.5 ventricles (Fig. 3B), consistent with previous studies ^6, 62^. We sought to determine whether defects in epicardial identity and activity associated with loss of WT1 ^6, 62^ impacted 6-*O*-endosulfatases. Within either the rare ventricular epicardial cells or the more abundant atrial population, *Sulf1* expression was strikingly downregulated whilst *Sulf2* was unexpectedly upregulated in *Wt1* KO hearts (Fig. 3C). These differences were more accurately quantified in E12.5 epicardial explants (Fig. 3D-G). We determined that the proportion of epicardial cells expressing medium to high levels of *Sulf1* (Bin 3 and Bin 4) dropped from 81% in controls to 57% in *Wt1* KO, accompanied by a 3.5-fold increase in epicardial cells with no or low detectable *Sulf1* transcripts (Bin 0 and Bin 1) compared to controls (Fig. 3G). In contrast, the proportion of epicardial cells expressing medium to high *Sulf2* transcripts (Bin 3 and Bin 4) increased from 21% in controls to 34% in *Wt1* KO, with a corresponding 26% decrease in epicardial cells that had no or low detectable *Sulf2* transcripts (Bin 0 and Bin 1) compared to controls (Fig. 3G).

**Fig. 3.**
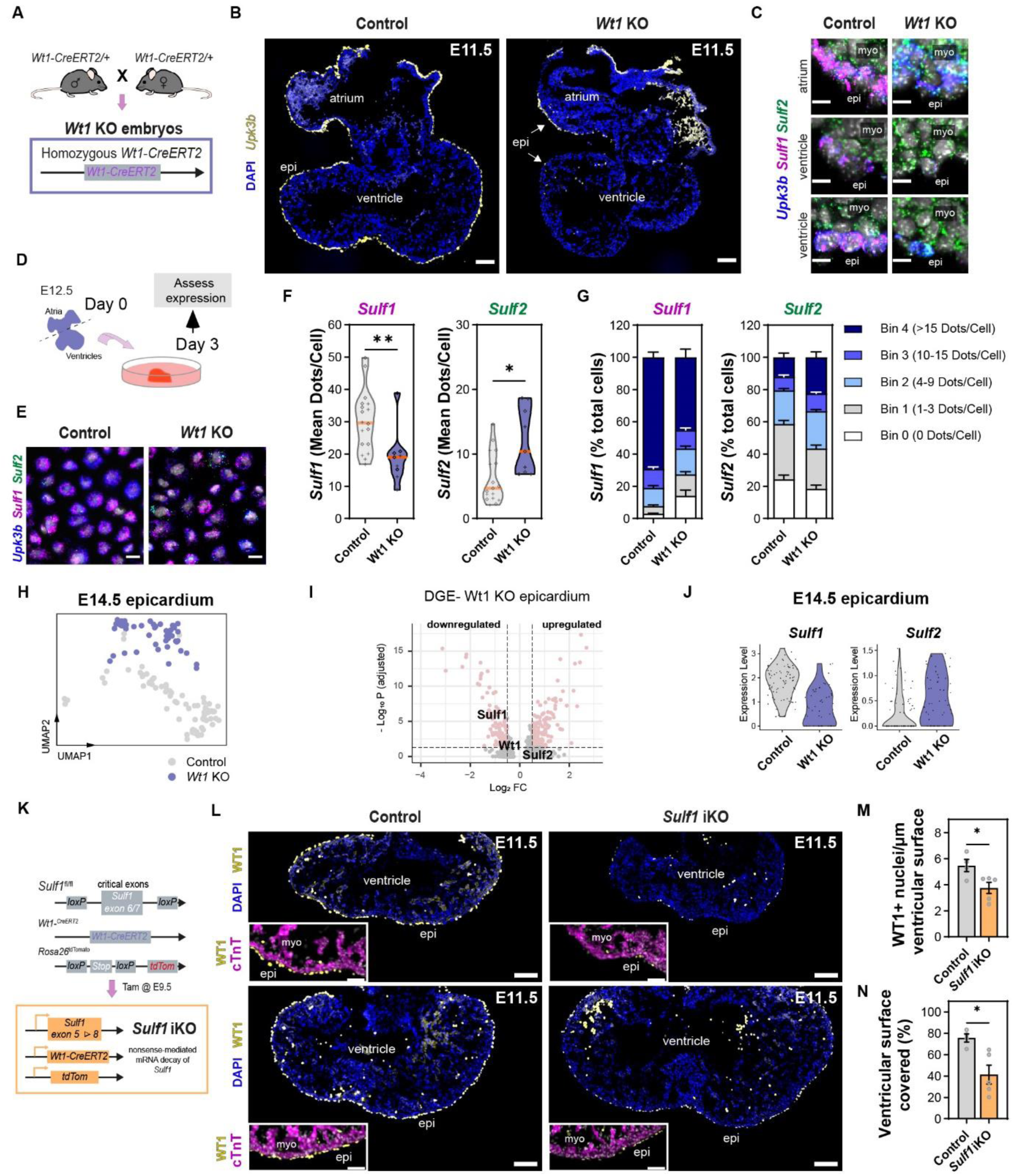
*Wt1* knock-out embryos demonstrate dysregulated 6-*O*-endosulfatase expression in the epicardium. (A) Generation of *Wt1* knock-out (KO) embryos by crossing heterozygous carriers of the *Wt1*-CreERT2 allele, an allele protein null for WT1. (B) Fluorescence ISH of E11.5 heart cryosections for *Upk3b* showing a severely disrupted ventricular epicardium in *Wt1* KO hearts. White arrows highlight epicardial cells. Scale bars: 100 µm. Images representative of n = 6 control and n = 5 *Wt1* KO hearts. (C) Fluorescence ISH of E11.5 heart cryosections for *Upk3b*, *Sulf1* and *Sulf2*, showing expression of *Sulf1* and *Sulf2* in the atrial and ventricular epicardium of control and *Wt1* KO hearts. Scale bars: 10 µm. Images representative of n = 6 control and n = 5 *Wt1* KO hearts. (D) Generation of epicardial explants to assess expression of *Sulf1* and *Sulf2* in control and *Wt1* KO hearts at E12.5. (E) Fluorescence ISH of control and *Wt1* KO epicardial explants for *Upk3b, Sulf1* and *Sulf2*, and corresponding quantification of (F) *Sulf1* and *Sulf2*. Scale bars, 20 µm. Images representative of n = 9-18 explants. Violin plot symbol cross (+) represents atrial explant; diamond (◊) represents ventricular explant. Orange line indicates the median (n = 9-18 explants). Unpaired t-test with Welch’s correction (*) P <0.05, (**) P <0.01. (G) Percentage of epicardial cells expressing *Sulf1* and *Sulf2,* grouped into 5 bins based on the number of dots (individual RNA molecules) per cell. Data are represented as mean ± SEM (n = 9-18 explants). (H) UMAP plot showing the epicardial cluster from control and *Wt1* KO embryonic mouse hearts at stage E14.5 (total 124 cells). (I) Volcano plot showing differential gene expression (DGE) in the *Wt1* KO epicardial cluster compared to control. Genes significantly upregulated or downregulated (filtering criteria adj_p<0.05, log2FC>0.5) coloured in pink. *Sulf1*, *Wt1* and *Sulf2* are annotated. (J) Violin plot showing relative expression of *Sulf1* and *Sulf2* in the control and *Wt1* KO epicardial cluster. (K) Generation of an inducible *Sulf1* knock-out mouse (*Sulf1* iKO) with lineage tracing capacity; restricted to the epicardial lineage (*Wt1*-CreERT2) and reported by tdTomato (Rosa26tdTomato). Successful recombination and removal of the critical region (KOMP) which includes exon 6 and 7 results in nonsense-mediated mRNA decay of the *Sulf1* transcript. (L) Immunofluorescence images showing WT1 positive epicardial cells on the surface of control and *Sulf1* iKO embryonic mouse hearts at E11.5. Inset shows selected region of interest of the right or left ventricular apex. Scale bar, 100µm and inset 50μm. Images representative of n = 4 control and n = 5 *Sulf1* iKO hearts. (M-N) Corresponding quantification of (M) WT1 positive epicardial cells and (N) coverage on the outer surface of the ventricular wall. *epi, epicardium; myo, myocardium*.

To determine whether altered *Sulf* expression persists to later stages of development, we analysed the published scRNA-seq dataset ^63^ of control and *Wt1* KO mice (*Wt1^GFPCre/GFPCre^*) at E14.5. In line with this study ^63^, the epicardial cluster separated according to mouse genotype (Fig. 3H) due to significant changes in the transcriptional profile, translating to defects in epicardial identity and activity. *Sulf1* was among the list of significantly downregulated genes in the *Wt1* KO epicardium, while *Sulf2* was upregulated, albeit not significantly (Fig. 3I, J). Overall, these findings show dysregulation of 6-*O*-endosulfases when the epicardium is impaired, with high *Sulf1* and limited *Sulf2* expression associated with normal epicardial formation and activity.

Given the abundance of *Sulf1* throughout the entire epicardium (Fig. 1 and Fig. 2) and its significant downregulation coincident with epicardial disruption (Fig. 3F, I), we questioned the role of *Sulf1* in epicardial formation and function. We generated inducible epicardium-restricted *Sulf1* knockout mice (*Sulf1* iKO; *Sulf1^fl/fl^*; *Wt1^CreERT2/+^*; Tamoxifen at E9.5), with *Rosa26^tdTomato^* lineage reporting (Fig. 3K). At E11.5, *Sulf1* iKO hearts displayed a striking disruption in ventricular epicardial formation (Fig. 3L), with a significant reduction in WT1 positive epicardial cells lining the ventricular surface (Fig. 3M), amounting to a 46% decrease in coverage (Fig. 3N). This demonstrates not only a requirement for *Sulf1* during epicardial formation, but also the remarkable extent to which *Sulf1* iKO embryos phenocopy *Wt1* KO, with respect to establishment of the epicardial layer. *Sulf1* iKO hearts were initially developmentally delayed, were typically smaller compared to controls, with reduced invasion of tdTomato-labelled EPDCs (Extended Data Fig. 4C) and often accompanied by myocardial compaction defects at E13.5 (Extended Data Fig. 4D, E). However, in contrast to the *Wt1* KO embryos which are embryonically lethal beyond stages E13.5-14.5 ^6, 63^, *Sulf1* iKO embryos were viable to at least E17.5 and showed similar composition of major cardiac cell types compared to controls (Extended Data Fig. 4F), likely due to the mosaic recombination achieved (Extended Data Fig. 4G). Although *Sulf1* was significantly downregulated in both the epicardium and its derivatives (Extended Data Fig. 4H), with levels halved in most cell types relative to controls (Extended Data Fig. 4I), ∼84% of epicardial cells still contained *Sulf1* transcripts (Extended Data Fig. 4I, J). Notwithstanding the remarkable capacity for non-targeted epicardial cells to compensate and replace the targeted population ^49^, functional compensation at multiple levels can be envisaged to permit normal HSPG-mediated signalling in the >80% of cells that retain residual *Sulf1* transcript. Taken together, despite the mosaicism and compensation, the transient phenotype shows all the hallmarks of previously reported epicardial defects ^6, 64, 65^, and suggests a functional role for Sulf1 in the developing epicardium.

### *Wt1* positively correlates with *Sulf1* in the epicardial lineage

To further explore the relationship between WT1 and 6-*O*-endosulfatases in the epicardial lineage, we assessed expression of these genes at E13.5, when the epicardium is at its most dynamic, undergoing proliferation and extensive EMT ^2^, coinciding with abundant production of factors to support myocardial and coronary vascular network development. Significant perturbation in these essential processes ^66^ commonly results in heart failure and embryonic lethality by this stage ^6, 63, 66, 67^. We analysed our E13.5 heart scRNA-seq data, subclustered the epicardial-lineage (tdTomato positive; Fig. 4A) and visualised the relative expression of *Wt1*, *Sulf1* and *Sulf2* in individual cells (Fig. 4B). The tdTomato-labelled lineage was composed of an equal proportion of epicardial cells and epicardium-derived mesenchymal cells, in line with previous reports ^2^. The epicardium-derived mesenchymal cells in our scRNA-seq dataset, are presumed to be the same tdTomato-labelled cells invading into the myocardium, enriched in the subepicardium and AVG at E13.5 (Extended Data Fig. 5A); later expected to contribute to cells of the ventricular myocardium, IVS and atrioventricular valve interstitium ^2^. *Wt1* and *Sulf1* strongly co-localised in the epicardium, and were consistently both downregulated in EPDCs (Fig. 4B) located in the subepicardium and myocardium (Fig. 4C, Extended Data Fig. 5A). Interestingly, *Sulf2* displayed an inverse correlation, being upregulated in EPDCs (Fig. 4B, Extended Data Fig. 5B).

**Fig. 4.**
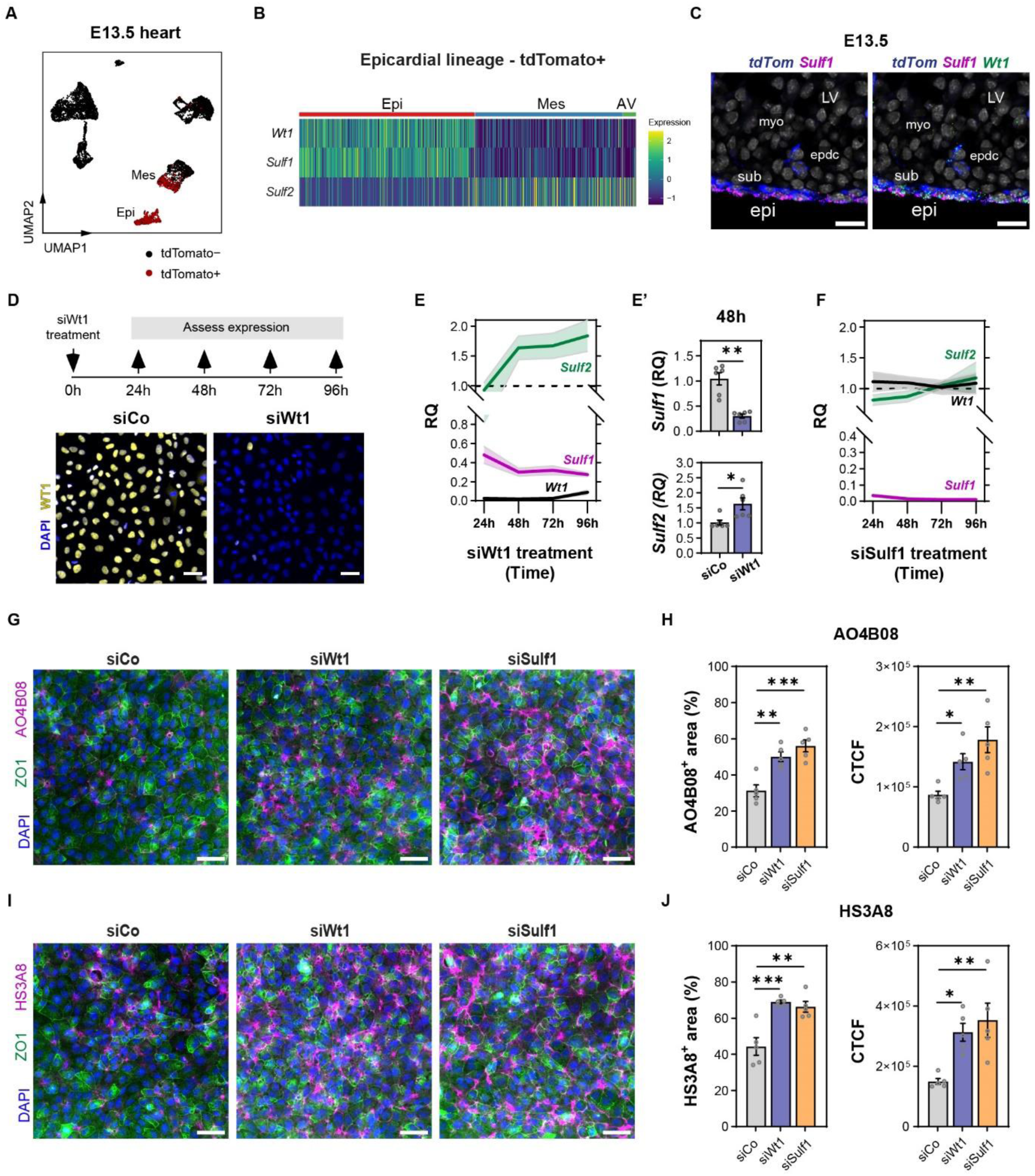
*Wt1* positively correlates with *Sulf1* expression in the epicardial lineage, and impacts HSPG sulfation. (A) UMAP plot showing *tdTomato* positive cells (epicardial lineage) in the E13.5 heart. (B) Heatmap showing expression of *Wt1*, *Sulf1* and *Sulf2* in the epicardial lineage subset (tdTomato+). High expression is indicated in yellow. (C) Fluorescence ISH of E13.5 heart cryosections for *tdTomato*, *Sulf1* and *Wt1*, showing downregulated expression of *Wt1* and *Sulf1* in epicardial-lineage subepicardial (sub) and mesenchymal cells invading the myocardium (epdc). Scale bars: 20 µm. Images representative of n = 3 hearts. (D) Mouse embryonic epicardial cell line (MEC1) transfected with control (siCo) or Wt1 siRNA (siWt1) for 24h, 48h, 72h and 96h. Immunofluorescence images of WT1 in control and Wt1 siRNA-transfected epicardial cells showing successful knock-down at 48h. Scale bar, 50 μm. (E) Time-course qRT-PCR analysis of *Wt1*, *Sulf1* and *Sulf2* relative expression in Wt1 siRNA-transfected epicardial cells. Values normalized to *Actb* and *Gapdh* and expressed relative to control siRNA treatment (dashed line), shaded data are represented as mean ± SEM (n = 4 - 6 experiments). (E’) Corresponding quantification of *Sulf1* and *Sulf2* relative expression in control and *Wt1* siRNA-transfected epicardial cells at 48h. Data are represented as mean ± SEM (n = 6 experiments). Unpaired t-test with Welch’s correction (*) P <0.05, (**) P <0.01. (F) Time-course qRT-PCR analysis of *Wt1*, *Sulf1* and *Sulf2* relative expression in Sulf1 siRNA-transfected epicardial cells. Values normalized to *Actb* and *Gapdh* and expressed relative to control siRNA treatment (dashed line), shaded data are represented as mean ± SEM (n = 4 - 6 experiments). (G) Immunofluorescence images showing anti-heparan sulfate staining (phage display antibody AO4B08) of epicardial cells transfected with siRNAs for 48h, and corresponding quantification of (H) percentage HS-positive area and corrected total cellular fluorescence (CTCF). Scale bars, 50µm. Images representative of n = 5 experiments. Data are represented as mean ± SEM (n = 5 experiments). One-Way ANOVA and Dunnett multiple comparison test, (*) P <0.05, (**) P <0.01, (***) P <0.001. (I) Immunofluorescence images showing anti-heparan sulfate staining (phage display antibody HS3A8) of epicardial cells transfected with siRNAs for 48h, and corresponding quantification of (J) percentage HS-positive area and corrected total cellular fluorescence (CTCF). Scale bars, 50µm. Images representative of n = 5 experiments. Data are represented as mean ± SEM (n = 5 experiments). One-Way ANOVA and Dunnett multiple comparison test, (*) P <0.05, (**) P <0.01, (***) P <0.001. *epi, epicardium; myo, myocardium; sub, subepicardium; epdc, epicardium-derived cells; LV, left ventricle*.

To further assess the dynamics of how WT1 changes 6-*O*-endosulfatases in the epicardial lineage, we utilised RNAi technology and the MEC1 mouse embryonic epicardial cell line, derived from E13.5 ventricular epicardial explants and shown to accurately reflect the state and plasticity of the ventricular epicardium ^68^. Temporal knock-down circumvents issues such as genetic mosaicism, compensation, phenotype penetrance and indirect regulation, and permits investigation of roles for WT1 and SULF1 in processes that occur beyond epicardial formation. We treated MEC1 epicardial cells with *Wt1* siRNA and achieved knock-down of ∼91% transcript and protein expression for up to 96h (Fig. 4D, E). In response to *Wt1* depletion, a 69% reduction in *Sulf1* expression was observed that plateaued at 48h (Fig. 4E, E’), while *Sulf2* expression demonstrated an increase of up to 1.7-fold by 96h (Fig. 4E), reaffirming the relationship between WT1 and 6-*O*-endosulfatase expression. To exclude the possibility of compensatory activity by the other isoform, we treated MEC1 epicardial cells with Sulf1 siRNA and detected no change in *Sulf2* expression (Fig. 4F). These results were in line with global *Sulf* knock-out studies ^40, 42, 44, 46^, and we concluded that *Sulf1* depletion did not result in upregulation of *Sulf2* in the epicardium. We used the same approach, to validate whether these differential changes occur in primary epicardial cells. After culturing E11.5 atria and ventricles over 24h to establish a layer of *Upk3b* and *Wt1*-expressing epicardial cells (Extended Data Fig. 5C), we treated explants with *Wt1* siRNA to achieve 35-53% transcript (proximal - distal location, respectively) and 87% protein knockdown at 48h (Extended Data Fig. 5D - F). E11.5 was selected as the epicardium is most proliferative ^69, 70^, guaranteeing adequate epicardial outgrowth within 24h and with limited spontaneous EMT after 3 days in culture. In accordance with the epicardial cell line experiments, we identified 36% reduction in *Sulf1* transcripts and 2-fold increase in *Sulf2* transcript levels in distally migrated cells (Extended Data Fig. 5F, G). It is worth noting we observed a change in the distribution of both *Wt1* and *Sulf1* transcripts with *Wt1* knock-down (Extended Data Fig. 5E), represented by predominantly nuclear-restricted mRNA clusters, as opposed to the abundant cytoplasmic puncta in controls (Extended Data Fig. 5H, I). Therefore, data in Extended Data Fig. 5F possibly underestimates the reduction in expression, further supported by a greater decrease identified at the protein level (Extended Data Fig. 5D). Taken together, these data implicate 6-*O*-endosulfatases during critical epicardial events, and presents these as potential gene targets for WT1 in the epicardium.

Extracellular SULF1 and SULF2 remove 6-*O*-sulfate moieties from HS chains ^38, 39^. Given *Wt1* depletion leads to downregulation of the predominant endosulfatase isoform, *Sulf1*, we sought to determine the impact on HSPG sulfation in epicardial cells. We analysed the 6-*O*-sulfate content of HSPGs in MEC1 epicardial cells, using phage display anti-HS antibodies, AO4B08 and HS3A8 ^71, 72^. The AO4B08 antibody which recognises both HS and heparin, with strong preference toward three consecutive *N*-, 6-*O*-sulfated disaccharide units ^73^, detected a significant increase in HS sulfation with Wt1 and Sulf1 siRNA treatment (Fig. 4G, H). The HS3A8 antibody which recognises highly sulfated domains within HS, with strong preference toward 6-*O*-, *N*-sulfated glucosamine ^71^, detected a significant increase in HS sulfation with both *Wt1* and *Sulf1* knockdown, compared to controls (Fig. 4I, J). Overall, these findings show *Sulf1* depletion in epicardial cells leads to reduced 6-*O*-endosulfatase activity characterised by an increase in sulfated HS chains, an effect that is replicated by reducing WT1 levels.

### WT1 binds intronic cis-regulatory element to enhance *Sulf1* transcription

The essential requirement for WT1 in cardiac development ^61^ has been associated, in part, with a proposed role in epicardial EMT ^6, 7^. The epithelial cell adhesion protein, E-cadherin (*Cdh1*), and its upstream regulator, Snai1, were identified in immortalised epicardial cells as gene targets of WT1 ^7^. However, *Cdh1* and *Snai1* were later challenged, when both were undetected in the embryonic epicardium, and epicardial EMT appeared to be independent of Snai1 ^6, 74^. In contrast, *bona fide* WT1 target genes in nephron progenitor cells have been identified during kidney development ^75^, and notably, 6-*O*-endosulfatases were amongst the list of targets identified in these cells ^75–77^. Unlike our findings in the epicardium, WT1 was associated with positive transcription of both 6-*O*-endosulfatases in kidney ^75, 76, 78^ and testis ^79^. In light of this, and the correlations observed in the epicardium, we sought to elucidate whether WT1 regulates 6-*O*-endosulfatase expression directly and how this may differ between *Sulf1* and *Sulf2*. We first assessed the landscape of chromatin accessibility in FACS sorted epicardial cells at E13.5 (CD31^−^tdTomato^+^PDPN^+^; Extended Data Fig. 6A, B), and employed TOBIAS footprinting ^80^ to identify WT1 binding to its candidate target sites in the *Sulf1* and *Sulf2* loci (Extended Data Fig. 6C, D). This analysis predicted WT1 binding to the highly accessible *Sulf1* promoter (Extended Data Fig. 6C), but no binding was predicted for *Sulf2* (Extended Data Fig. 6D). As expected, there was lower accessibility in this locus which reflected reduced *Sulf2* gene activity in the epicardial layer. Subsequent CUT&RUN-seq experiments that mapped genome-wide binding sites of WT1 in epicardial cells (MEC1; Fig. 5A) confirmed weak, although variable across replicates, binding of WT1 to the *Sulf1* promoter (ROI I, Fig. 5B). In contrast, our analysis revealed significant WT1 binding to an intronic enhancer (E1), marked by active chromatin modification H3K27ac, located in intron 1 of the *Sulf1* gene (ROI II, Fig. 5B). This region was accessible in freshly isolated E13.5 epicardial cells and contained two WT1 binding sites annotated from the JASPAR database (Extended Data Fig. 6E). Next, we applied the Activity-By-Contact (ABC) model ^81^ incorporating CUT&RUN-seq for histone marks, ATAC- and RNA-seq data, and we identified a significant interaction between intronic enhancer E1 and the upstream *Sulf1* promoter (Extended Data Fig. 6E). Interestingly, the CUT&RUN-seq experiments also revealed significant binding of WT1 to the third intron of the *Sulf2* gene (Fig. 5C), displaying chromatin accessibility (Extended Data Fig. 6D) without any histone marks or gene activity (ROI II, Fig. 5C).

**Fig. 5.**
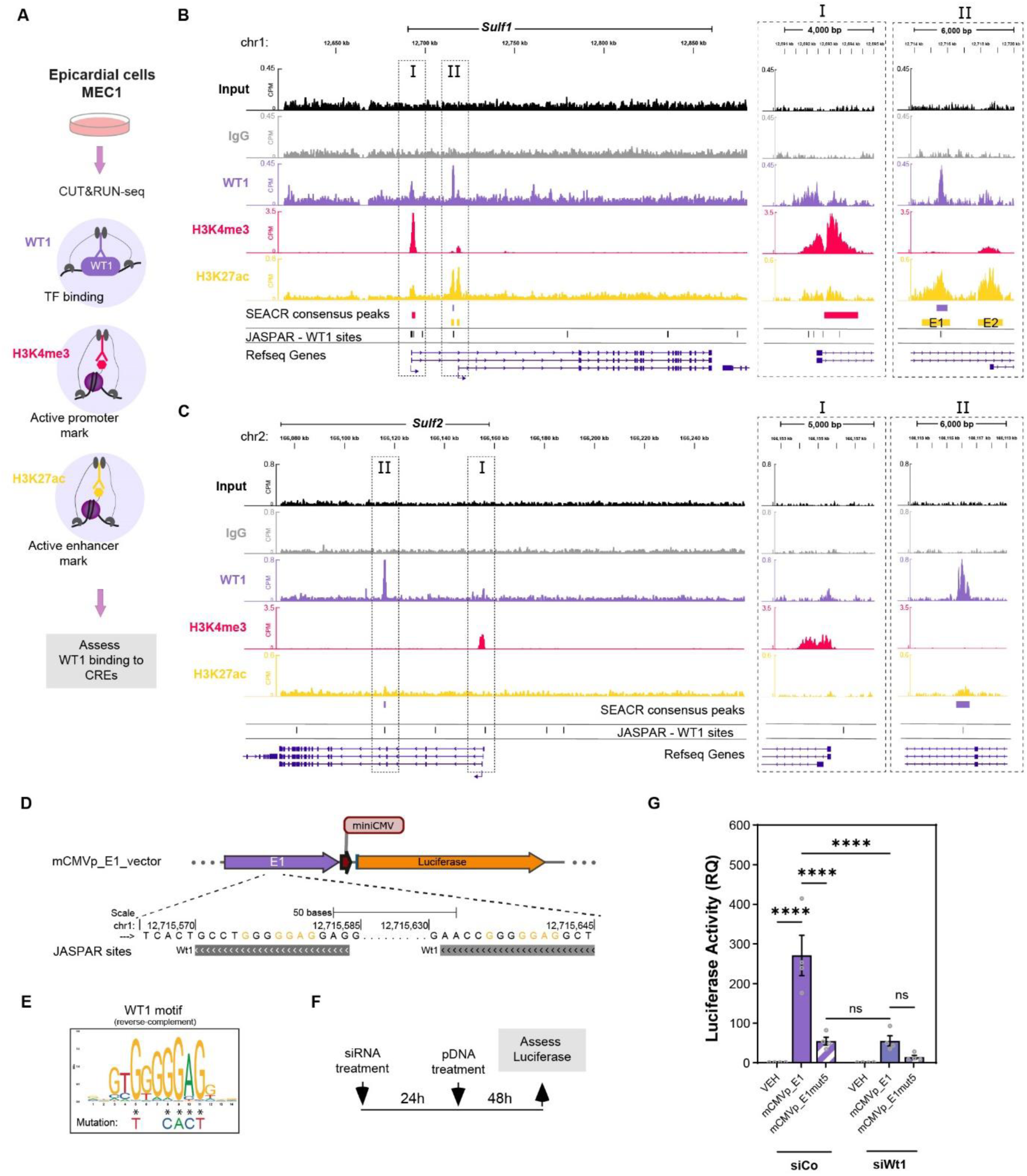
WT1 binds an intronic enhancer proximal to the *Sulf1* promoter. (A) CUT&RUN-seq of the epicardial cell line (MEC1) to assess WT1 transcription factor binding and histone marks. (B-C) Genome tracks showing binding of WT1 at the (B) *Sulf1* and (C) *Sulf2* locus. Track signal is in counts per million (CPM) and merged from n = 2 independent replicates. SEACR consensus peaks calculated from n = 3 (WT1 and H3K27ac) and n = 4 (H3K4me3 and IgG) independent replicates. Region of interest (ROI) indicated in inset I, promoter/transcription start site; inset II, intronic WT1 binding. JASPAR database WT1 binding sites indicated. (D) Luciferase reporter vector containing intronic enhancer-*Sulf1* fragment E1 upstream of a minimal CMV promoter and Luciferase gene (mCMVp_E1). Two WT1 binding sites located within the E1 fragment sequence. (E) 5 nucleotide mutation of the WT1 motif located within the enhancer-*Sulf1* fragment E1 (mCMVp_E1mut5) to decrease WT1 binding affinity. (F) MEC1 epicardial cell line transfected with control (siCo) or Wt1 siRNA (siWt1) for 24h, followed by transfection with mCMVp_E1 or mCMVp_E1mut5 for 48h. Corresponding quantification of (G) Luciferase activity to assess transcriptional activity of WT1 on intronic *Sulf1* enhancer in the epicardial cell line. Values expressed relative to vehicle control. Data are represented as mean ± SEM (n = 4 experiments). One-Way ANOVA and Šídák multiple comparison test (****) P <0.0001 and ns, not significant.

To validate the functional role of WT1 binding to the intronic enhancer E1 within the *Sulf1* gene, we performed reporter assays in cultured MEC1 epicardial cells, with or without WT1 present (Fig. 5D-G, Extended Data Fig. 6F). We cloned the intronic enhancer sequence into a luciferase reporter vector containing a miniCMV promoter and the luciferase gene (mCMVp E1, Fig. 5D). We also generated an alternative vector (mCMVp E1mut5) incorporating a five-nucleotide mutation in both WT1 motif sites (Fig. 5E), predicted to abolish WT1 binding without introducing alternative motifs recognised by transcription factors expressed in epicardial cells. Transfection of epicardial cells with mCMVp E1 demonstrated significant luciferase activity, which was attenuated with mutation of the WT1 binding sites, or with knockdown of WT1 using siRNA (Fig. 5G, Extended Data Fig. 6F). These data indicate dependency of enhancer activity on WT1 binding, and proposes that this regulatory element is responsible for transcriptional control of *Sulf1*.

### Epicardial cells are primary orchestrators and responders in HSPG-dependent signalling

Having demonstrated transcriptional regulation of *Sulf1* by WT1, we next asked whether this regulatory relationship impacts specific HSPG-dependent pathways in the E13.5 epicardium. In order to infer global HSPG-dependent signalling occurring in the E13.5 heart, we used scRNA-seq and applied CellChat ^82^ with an updated database, which we supplemented with literature-supported ligand-receptor interactions from CellPhoneDB ^83^, scTalkDB ^84^ and STRINGDB (experimental interaction), and further annotated pathways associated with HSPG molecules. The epicardium demonstrated the most incoming and outgoing HSPG-dependent signalling interactions, closely followed by the endocardium (Fig. 6A). We focused our attention on incoming signalling to the epicardium, and identified a broad range of HSPG-dependent pathways (Fig. 6B), most of which originated from epicardial, mesenchymal and endocardial senders (Fig. 6C).

**Fig. 6.**
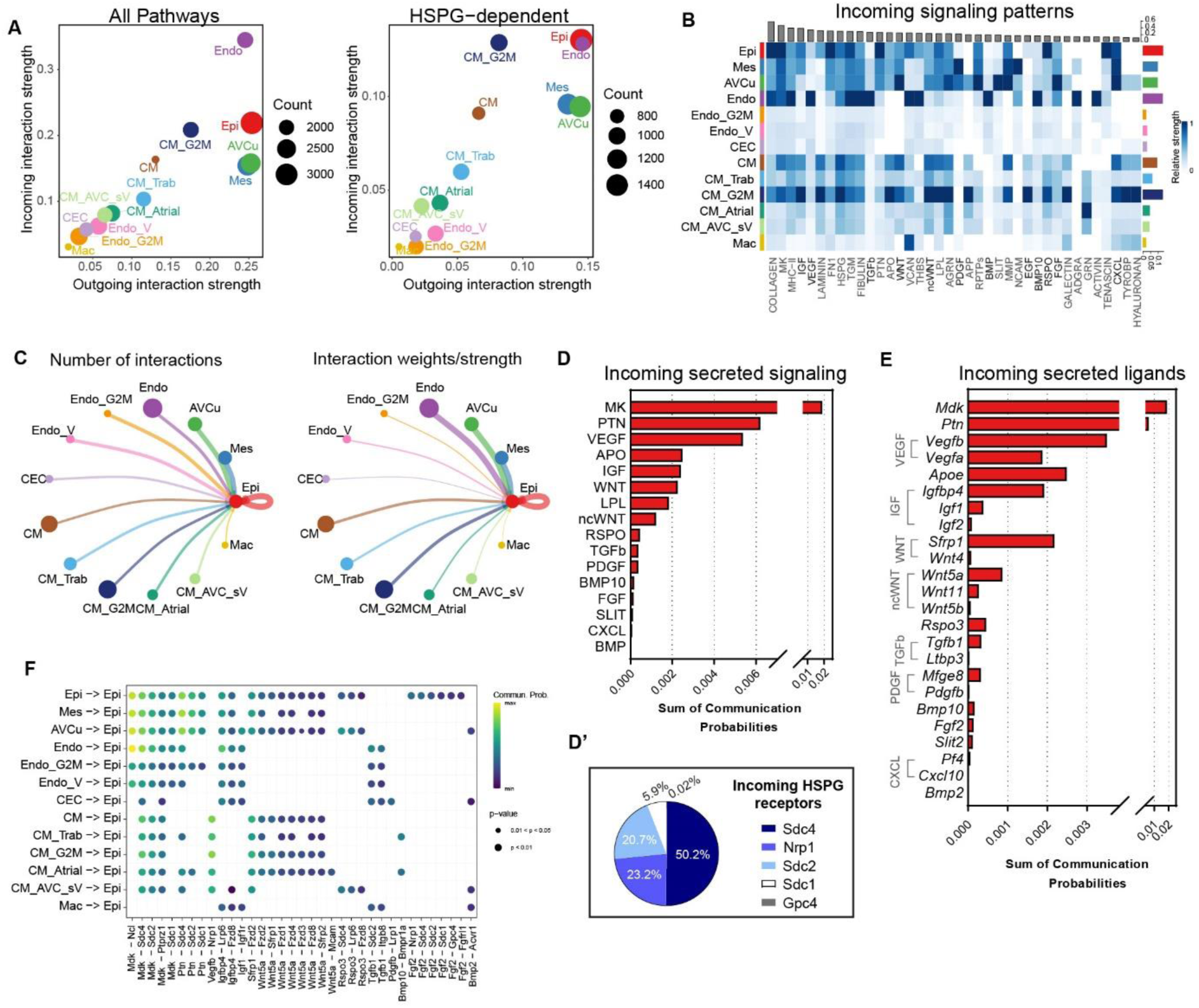
HSPG-dependent pathways identified in the E13.5 epicardium. (A) Cell-cell interaction strengths plotted for all cell types, divided into total (All pathways) and HSPG-dependent pathways, indicating incoming and outgoing interactions. (B) Heatmap and associated bar plots showing most incoming HSPG-dependent signalling patterns. Heatmap represents relative signalling strength of pathway across cell types. Coloured bar plot shows the total signalling strength of a cell type by summarizing all signalling pathways. Grey bar plot shows the total signalling strength of a signalling pathway by summarizing all cell types. High communication probability is indicated in dark blue. (C) Circle plots of incoming HSPG-dependent signalling networks to the epicardium. (D) Incoming secreted signalling received by the epicardium ranked from highest to lowest communication probabilities. (D’) Pie chart showing percentage contribution of HSPG molecules as receptors on the epicardium. (E) Incoming secreted ligands received by the epicardium ranked from highest to lowest communication probabilities. (F) Dot plot indicating source cell types communicating with the epicardium and corresponding key ligand-receptor pairs. High communication probability is indicated in yellow, dot size indicates p-values. *AVCu, atrioventricular cushion; CEC, coronary endothelial cells; CM, ventricular cardiomyocytes; CM_Atrial, atrial cardiomyocytes; CM_AVC_sV, atrioventricular canal and sinus venosus cardiomyocytes; CM_Trab, trabecular cardiomyocytes; Endo, endocardial cells; Endo_V, valvular endothelial and endocardial cells; Epi, epicardium; G2M, proliferating; Mac, macrophages; Mes, mesenchyme*.

Next, we ranked significant incoming signalling (secretory factors) according to the sum of communication probabilities and identified shared pathways involving MDK, VEGF, WNT family members and TGFβ in the top ten (Fig. 6D), some of which were shown to signal in the embryonic epicardium ^6, 13, 16, 85^, while others are currently unexplored, or shown to be transmitted, rather than received, by the epicardial lineage ^86–92^. In our analysis, we found that SDC4 and NRP1 – expressed on epicardial cells – contributed the majority of HSPG receptor interactions, with SDC4 participating in 50.2% of incoming ligand-HSPG communications (Fig. 6D’). To prioritise candidate factors for functional validation, we separated pathways into their participant ligands (Fig. 6E), and selected factors according to their communication probability scoring and HSPG receptors (Fig. 6F, Extended Data Fig. 7A). We were also interested in interrogating factors that formed cell type exclusive interactions, such as VEGFB and BMP10 incoming from cardiomyocytes, PDGFB from CECs and FGF2 from epicardial cells (Fig. 6F). Our findings suggest the epicardium is subjected to significant HSPG-dependent signalling at E13.5, mapping a complex communication potential, with SDC4 as principal HSPG partner, and revealing both recognised and new factors that may coordinate cellular responses in this tissue.

### WT1 regulation of SULF1 activity modulates epicardial proliferation and EMT

We interrogated twelve factors shortlisted from our ligand-receptor pair analysis based on properties described above, along with TGFβ2 as a positive control, since it was previously shown to activate the MEC1 line ^19, 93^. We first questioned how these factors influence epicardial proliferation and EMT, and assessed their activity in the presence or absence of WT1 and SULF1. We treated MEC1 epicardial cells with *Wt1* or *Sulf1* siRNA, to reduce SULF1 activity, before treating with selected factors and assessing proliferation rate using EdU incorporation (Fig. 7A). Only FGF2 promoted epicardial proliferation (1.3-fold increase), while TGFβ and BMP10 decreased proliferation by 75% and 38%, respectively (Extended Data Fig. 7B). We noted that *Wt1* and *Sulf1* knockdown alone reduced EdU positive cells by 35-37%, however, both demonstrated a significant increase in the proliferation rate upon FGF2 stimulation, compared with control transfection (Fig. 7B, C). Taken together, *Sulf1* depletion elicits a stronger proliferative response to FGF2 in epicardial cells, verifying a HSPG-dependent pathway impacted by the WT1-*Sulf1* regulatory relationship.

**Fig. 7.**
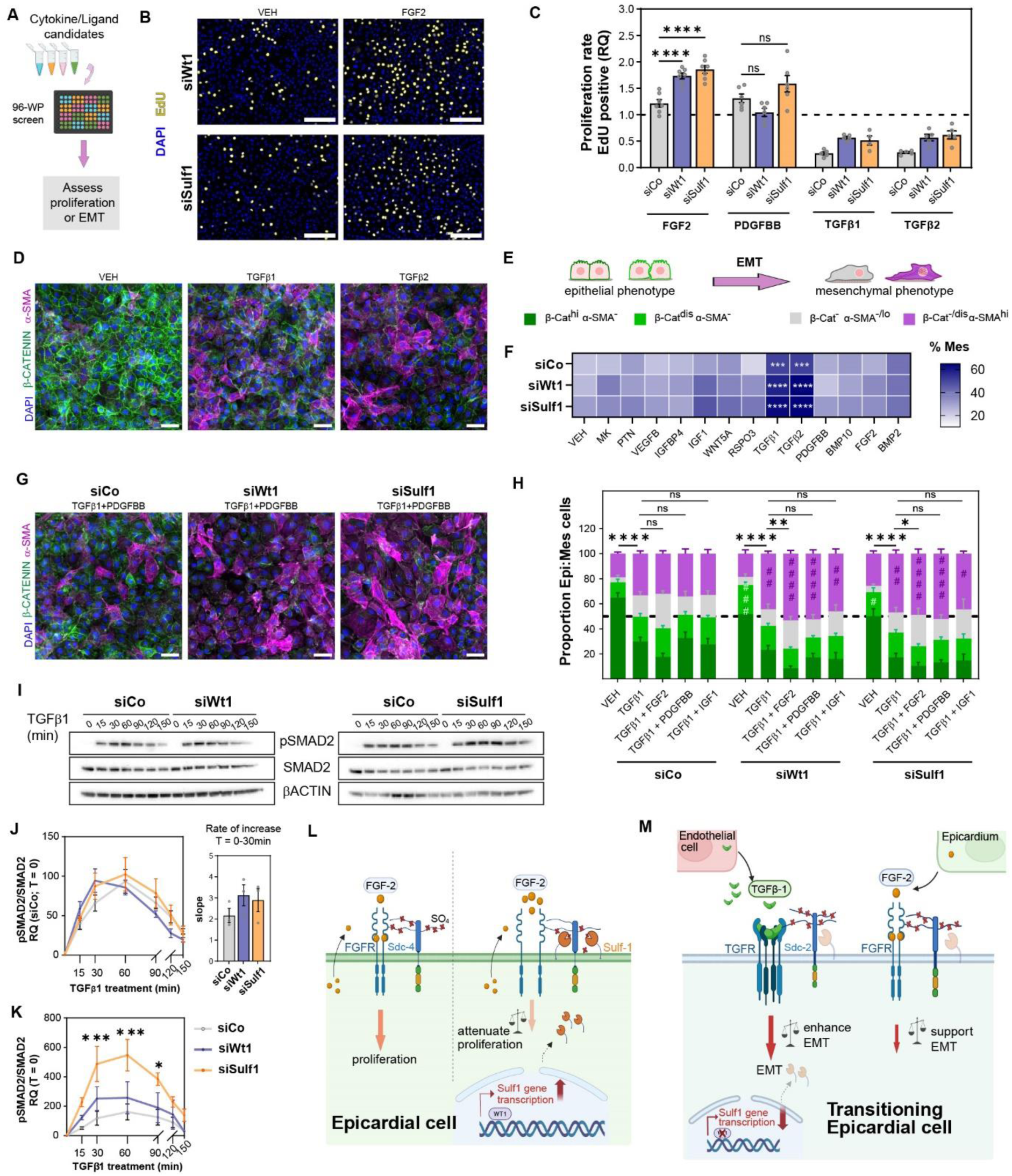
WT1 regulation of Sulf1 activity impacts HSPG-dependent pathways modulating epicardial proliferation and EMT. (A) MEC1 epicardial cell line with knock-down of specific target genes, were treated with candidate cytokines to assess proliferation and epithelial-to-mesenchymal transition (EMT). (B-C) Immunofluorescence images showing EdU positive epicardial cells transfected with control (siCo), Wt1 (siWt1) or Sulf1 siRNA (siSulf1) for 48h, followed by treatment with candidate cytokines for 24h, and corresponding quantification of the (C) rate of proliferation. Scale bars, 200µm. Images representative of n = 11 experiments. Values expressed relative to their vehicle control. Data are represented as mean ± SEM (n = 4 – 7 experiments). One-Way ANOVA and Šídák multiple comparison test, (****) P <0.0001 and ns, not significant. (D-F) Immunofluorescence images showing β-CATENIN and α-SMA staining of epicardial cells transfected with siRNAs for 48h, followed by treatment with candidate cytokines for 24h, and corresponding quantification of (F) percentage mesenchymal cells. Scale bars, 50µm. Images representative of n = 9 experiments. Data expressed as mean (n = 4 – 9 experiments). Two-Way ANOVA and Dunnett multiple comparison test, (***) P <0.001, (****) P <0.0001. (G-H) Immunofluorescence images showing β-CATENIN and α-SMA staining of epicardial cells transfected with siRNAs for 48h, followed by treatment with candidate cytokines for 24h, and corresponding quantification of (H) percentage epicardial and mesenchymal cells. Scale bars, 50µm. Images representative of n = 8 experiments. Data are represented as mean ± SEM (n = 7 – 9 experiments). One-Way ANOVA and Fisher’s LSD test (*, ^#^) P <0.05, (**, ^##^) P <0.01, (***, ^###^) P <0.001, (****, ^####^) P <0.0001 and ns, not significant. Colour of bars and hash (#) symbol represent cell phenotype. Asterisk symbols represent mesenchymal cell comparison between treatment groups, and hash symbols represent comparison between their respective siControl group (e.g. siControl + TGFβ1 vs siWt1 + TGFβ1) for a specific cell phenotype. (I) Western blot showing SMAD2 signalling in control (siCo), Wt1 (siWt1) and Sulf1 (siSulf1) siRNA-transfected epicardial cell line (48h), followed by treatment with 10ng/ml TGFβ1 (time response). Blot representative of n = 3 experiments. (J-K) Corresponding quantification of SMAD2 signalling expressed relative to (J) control siRNA at time = 0, and (K) corresponding siRNA at time = 0. Data are represented as mean ± SEM (n = 3 experiments). Two-Way ANOVA and Dunnett multiple comparison test (*) P <0.05, (***) P <0.001. (L) Model of transcription factor WT1 positively regulating *Sulf1* expression to temporarily attenuate autocrine FGF2-stimulated proliferation in the epicardium. (M) Model of WT1 downregulation directly downregulates *Sulf1* expression in transitioning epicardial cells to promote a robust response to EMT signals coming from endothelial cells, with the EMT process supported by the adjacent FGF2-expressing epicardium. Image in L and M created with BioRender.com

We used a similar approach to assess impact on epicardial cell transition and differentiation, by screening the same twelve factors and quantifying mesenchymal derivatives. We immunostained MEC1 epicardial cells with β-CATENIN, an intracellular component of anchoring junctions that enable interaction with neighbouring cells ^94^, and a classic hallmark of epithelial monolayers. We co-stained for α-smooth muscle actin (α-SMA), to label actin filaments, characteristic of cells adopting a mesenchymal phenotype. Morphological changes consistent with adoption of a mesenchymal phenotype were hallmarked by disruption or complete loss of cell-cell junctions (β-CATENIN negative or translocated) or upregulated α-SMA with spindle-shaped morphology (Fig. 7D, E). TGFβ treatment significantly increased the percentage of mesenchymal cells with 50% in the control siRNA supplemented with TGFβ, which was augmented to 58% and 63% with *Wt1* and *Sulf1* knockdown, respectively (Fig. 7F, Extended Data Fig. 7C). Epicardial cells with reduced SULF1 activity also demonstrated a significant increase in EMT with IGF1 treatment (Extended Data Fig. 7C). Considering other factors present in the heart could influence cell fate decision once EMT is induced, we next examined TGFβ1 in combination with other shortlisted HSPG-dependent factors (Fig. 7G, H), and found that FGF2 expands the proportion of mesenchymal cells displaying complete loss of β-CATENIN without α-SMA upregulation (grey; Fig. 7H). Interestingly, we detected a 1.9-fold and 1.6-fold increase, respectively, in the proportion of epicardial cells with disrupted cell-cell contacts (light green) in response to *Wt1* and *Sulf1* knockdown alone (Fig. 7H). Disruption in epicardial cell-cell contacts was previously reported following *Wt1* knockdown in epicardial explants derived from hearts at E12.5 ^95^. Further, *Wt1* knockdown in human adult epicardial cells demonstrated morphological changes that suggested greater spontaneous EMT *in vitro* ^96^. These data consistently support a role for WT1 in maintaining epithelial state, and implicate Sulf1 as a key downstream effector. Both *Wt1* and *Sulf1* knockdown enhanced the proportion of α-SMA-high mesenchymal cells following FGF2, PDGFBB, and IGF1 combinatorial treatments, with the greatest increase in EMT associated with FGF2/TGFβ in combination (Fig. 7H). We concluded that *Sulf1* depletion elicits a robust response to TGFβ in epicardial cells, enhancing EMT, and further permitting FGF2 to aid in the process.

We validated the expression domains of *Tgfb1, Tgfb2* and *Fgf2* by fluorescent ISH staining and assessed the positioning of signalling cells in the E13.5 heart (Extended Data Fig. 7D - F), particularly with regard to their proximity to the epicardium and EPDCs. scRNA-seq analysis showed that ECs and macrophages express *Tgfb1* (Extended Data Fig. 7D), and we further validated abundant expression of this factor in the endocardium and subepicardial domain of E13.5 hearts (Extended Data Fig. 7D’). Indeed, our ligand-receptor inference analysis highlighted that most significant TGFβ1 communications to the epicardium (Fig. 6F, Extended Data Fig. 7A) originated from the non-proliferative endocardial cluster and CECs. *Pecam1* positive CECs sprouting down from the sinus venosus at the base of the heart typically run along the subepicardial space (Extended Data Fig. 7D’) before invading into the compact myocardium ^97^. Our data presents proximity to the tdTomato-labelled epicardium and derivative mesenchymal cells, even showing side-by-side alignment with *Tgfb1*-expressing CECs in the interventricular septum (Extended Data Fig. 7D’). In contrast, we found broad *Tgfb2* expression throughout cardiac cell types (Extended Data Fig. 7E). We confirmed abundant expression in the epicardium, cardiomyocytes and mesenchyme of the atrioventricular groove and cushion. Interestingly, expression of *Tgfb2* in regions with high EMT activity was mainly contributed by the tdTomato-labelled epicardium-derived mesenchymal cells themselves (right inset, Extended Data Fig. 7E’). Considering the abundance of *Tgfb2*, we asked why this known inducer of EMT ^14, 19, 93^ was not predicted in our analysis of HSPG-dependent pathways. We identified that although ∼20-37% of the E13.5 epicardium expresses the requisite TGFβ receptors 1 and 2 (Extended Data Fig. 7G), it does not express the HSPG co-receptor *Tgfbr3* (Extended Data Fig. 2A) essential for TGFβ2 binding to the receptors ^98^. In contrast, *Tgfbr3* is expressed in cultured epicardial cells, supported by the enrichment of H3K4me3 at the *Tgfbr3* promoter (Extended Data Fig. 7H) and previously reported gene expression ^19, 99^ in MEC1 cells, which may explain their ability to respond to this factor *in vitro*. We concluded that TGFβ1, as opposed to TGFβ2, expressed by ECs is more likely to promote EMT and support the transition of derivative mesenchymal cells. We further identified that the epicardium is the major source of *Fgf2* (Extended Data Fig. 7F, F’), suggesting autocrine regulation of its proliferation. Taken together, epicardial-expressed SULF1 is important in both paracrine and autocrine signalling cues which regulate proliferation and contribution to the developing heart.

While both FGF2 ^35, 100^ and TGFβ1 ^101, 102^ directly bind HS, the mechanism by which HS 6-*O* regulates their signalling differs. Nonetheless, 6-*O*-sulfation was required for FGF2-FGFR1 signalling in MEFs and HUVECs ^35, 103^, and for TGFβ1 signalling in lung fibroblasts ^104^. Lastly, we interrogated how transcriptional regulation of *Sulf1* by WT1 may impact the downstream HSPG-dependent signalling response, to bring about the observed functional changes in epicardial cells, that suggest a mechanism of growth factor sensitisation. We examined TGFβ1 signalling in epicardial cells and found that reduced SULF1 activity, achieved by either *Wt1* or *Sulf1* knockdown, enhanced SMAD2 phosphorylation (Fig. 7I). Intriguingly, our analysis showed that both the rate of increase in the first 30 minutes of TGFβ1 treatment (Fig. 7J, all relative to siControl at T=0) and overall magnitude of phospho-SMAD2 signalling (Fig. 7K, relative to their respective siRNA treatment at T=0) was enhanced in knockdown conditions. These data support enhanced TGFβ1 signalling with reduced SULF1 activity, and suggest the WT1-*Sulf1* regulatory interaction may influence epicardial cell response to EMT-inducing factors.

## Discussion

Impairing epicardial function, by genetic deletion of key transcription factors ^6, 63, 105^ and essential signalling ^66, 89, 106–108^, or by direct ablation ^109–111^, proves detrimental to both heart development and repair in disease. Despite improvements in our understanding of embryonic epicardial properties, little is known about how the epicardium deciphers signalling cues to temporally and spatially orchestrate cellular processes. In this study, we explored 6-*O*-endosulfatases capable of modifying the ‘sugar code’ extracellularly and uncovered a transcriptional regulatory relationship between the critical epicardial regulator WT1 and 6-*O*-endosulfatase isoform Sulf1. We propose that this relationship instructs changes in HS sulfation, to adjust epicardial lineage sensitivity to signalling cues, and to modulate cellular responses during heart development (Figure 7L, M).

Before exploring this regulatory relationship in detail, we first delineated *Sulf1* and *Sulf2* cell type-specific expression patterns, which along with the HSPG diversity we observed in the embryonic heart, reinforces the notion of temporal and regional HS patterning in organs dynamically changing over the course of development ^38, 39^. Here, we recognised cardiomyocytes as the major source of Sulf2, while an abundant and localised expression of Sulf1 was detected in the epicardium. Our finding that 6-*O*-endosulfatase isoforms show distinct expression profiles in the embryonic heart, with only a minor overlap, supports their functional diversity *in vivo* ^40, 45, 112^. Embryonic lung, gonads and eyes similarly show non-overlapping expression domains ^41, 113^. Notably, enriched *Sulf1* expression in the epicardium builds upon evidence that this isoform is commonly found in embryonic mesothelia, epithelia and connective tissue lining organs, with functions akin to the epicardium during development. Examples include lung pleura ^41, 43, 113^, pericardium ^41, 113^, pancreatic mesothelium ^113^ and periosteum ^113^. Conservation between species is likely as *Sulf1* homologs are enriched in the *Xenopus* pericardium ^114^ and human epicardium ^115^. *Sulf1* transcription in visceral mesothelia, such as that of lung and pancreas, may also be subject to control by WT1 during embryogenesis ^116–118^. Thus, the modulation of cellular processes by the WT1-*Sulf1* regulatory relationship that we describe here may apply more broadly across other organs and species.

Our findings that loss of WT1 in epicardial cells confers an incomplete loss of *Sulf1* supports the notion that WT1 acts by enhancing *Sulf1* expression, and aligns with other reports in mesonephros and podocytes ^76, 78, 119^. Our data offer improved mapping of WT1-DNA binding sites in the *Sulf1* locus, as well as across the genome. When paired with histone mark information, we were able to identify that, in the case of *Sulf1*, WT1 transcriptional regulatory activity is exerted through direct binding to an intronic enhancer, a region interacting with the promoter. Intriguingly, a similar region was identified in adult glomeruli with high placental mammal conservation ^119^. In this study, we uncovered notable differential expression of 6-*O*-endosulfatase isoforms between the epicardium and its early mesenchymal derivatives. Specifically, *Sulf1* was downregulated while *Sulf2* was upregulated at the time of transition. We went on to expand the relationship between *Wt1* and *Sulf1* following EMT, finding it comparable to human foetal hearts, where both genes downregulate in epicardium-derived mesenchymal cells ^115^. In contrast, immature podocytes and adult sertoli cells feature continued activation of both 6-*O*-endosulfatase isoforms by direct binding of WT1 to their promoters ^75, 76, 79^. We propose that *Sulf2* takes on a lesser role in the epicardium, as fewer epicardial cells express this isoform and do so at lower levels. Unlike *Sulf1* transcription, expression appears independent of key regulator WT1. Rather, an inverse relationship exists between *Wt1* and *Sulf2* in the epicardial lineage, which we speculate results from adoption of mesenchymal identity. Insights from cancer research may be relevant, as *Sulf2* is frequently associated with the terminal mesenchymal state in EMT and, as such, underpins the invasive nature of these cells in tumour metastasis ^120, 121^.

Despite challenges in studying HS-modifying enzymes *in vivo* ^39, 41, 42, 44, 45, 122^, we identified a defect in the formation of the epicardium with *Sulf1* deficiency, likely due to dysregulated BMP, FGF and WNT signalling during early epicardial specification ^8, 12, 123^. While SULF1 activity may limit FGF, it is required for BMP and WNT signalling ^124, 125^. Further, the impaired epicardial phenotype we observed in *Sulf1* iKO hearts partly mimicked the phenotype reported for *Wt1* nulls ^6, 62, 63^, strengthening functional association between these two genes. In clinical cases where mutations in *WT1* underlie inherited disease, dysregulated cellular processes during kidney development associate with downregulation of *SULF1* in patient cells ^126^. This raises questions about whether congenital heart defects associated with *WT1* mutations ^127–129^, although rare, involve disrupted SULF1 activity in the epicardial lineage.

Here, we identified dynamic epicardial *Sulf1* expression, with peak levels at E13.5 coinciding with shifts in environmental signals and cellular processes. In line with others ^69, 70^, we identify that the E13.5 epicardium is less proliferative overall and propose that Sulf1 temporarily limits autocrine FGF2-stimulated proliferation. In *Drosophila*, Sulf1 ensures termination of intestinal stem cell division and return to homeostasis, as it limits mitogenic pathways initially required in the regenerative response ^130^. An enhanced mitogenic response to FGF2 upon *Sulf1* downregulation was reported in embryonic and cancer cells ^39, 42, 122, 131, 132^, and supports our findings of sensitisation in *Sulf1*-depleted epicardial cells.

6-*O-*endosulfatases have been implicated in regulating cell fate in a variety of stem and progenitor cell niches ^43, 44, 133–135^. We and others have identified that the epicardium undergoes EMT over a limited developmental window, with apparent regionalised ‘waves’ of invasion, first appearing at the atrioventricular grooves around E11.5 and becoming more widespread by E13.5 ^2, 136–138^. Here, we identify that TGFβ1 signalling in the epicardium is impacted by the WT1-*Sulf1* regulatory relationship. We show that over a third of the epicardial population expresses TGFβ receptors required to respond to the distinct sources of TGFβ1 *in vivo*. We propose that the endocardium forms a gradient of TGFβ1 in the ventricular myocardium, and transient downregulation of *Sulf1* in EPDCs promotes a robust response to EMT signals, with their transition supported by the adjacent TGFβ1/PDGFB-expressing coronary endothelium and FGF2-expressing epicardium. With downregulation of WT1 lying upstream of this process, the *Sulf1*-low epicardial lineage thus becomes more sensitised to EMT factors, leading to elevated intracellular signalling. Indeed, increased TGFβ1 signalling was previously reported in *Sulf1*-deficient lung fibroblasts, allowing for enhanced transdifferentiation to myofibroblasts ^139^. Curiously, at the final stages of cardiac development, epicardium-derived mesenchymal cells situated deeper within the myocardium, compared to those more superficially, expressed higher levels of *Sulf1*, possibly a consequence of establishing fibroblast cell fate ^2, 46, 140^.

The membrane-bound localisation of 6-*O*-endosulfatases urges us to recognise that their impact on HSPG-dependent signalling is not restricted to autoregulatory epicardial processes. Our identification of the epicardium as the major source of outgoing HSPG-dependent signalling and SULF1 activity raises questions as to how this may influence paracrine signalling to diverse adjacent cell types. For example, SULF1 activity was suggested to enhance the angiogenic effects of CXCL12 ^46^, a factor secreted by the epicardium and important in coronary arteriogenesis during development ^60, 141^.

The epicardium seems to lose its plasticity in adulthood, and is considered largely quiescent under physiological conditions. Although in injured hearts upregulated *Sulf1* expression is largely contributed by cells within the infarct zone ^46, 142^, it has also been detected in the reactivated ventricular epicardium ^47^. Interestingly, *Sulf1* levels downregulate at 3 to 6 days post-myocardial infarction injury as the epicardium undergoes EMT to derive the subepicardial mesenchyme ^47^. We speculate that this change in transcription may arise from temporal activation of *Wt1* ^5, 143, 144^, which raises questions as to whether the regulatory relationship we describe in development may be recapitulated in cardiac injury.

Overall, our work provides evidence of transcriptional regulation of a HS-modifying enzyme in the developing epicardium, to modulate cellular processes critical to heart formation and function. Whether this regulatory relationship additionally impacts paracrine HSPG-dependent signalling in epicardial events following cardiac injury remains to be explored.

## Methods

### Animal models

All procedures were approved by the University of Oxford Animal Welfare and Ethical Review Board, in accordance with Animals (Scientific Procedures) Act 1986 (Home Office, UK). Mice were maintained in individually ventilated cages (IVCs) and ventilated racks at 22 °C and 55% humidity. Mouse strains used include C57BL/6J (Charles River), Gt(ROSA)26Sor^tm14(CAG-tdTomato)Hze^/J (MGI ID: 3809524) or Gt(ROSA)26Sor^tm9(CAG-tdTomato)Hze^/J (MGI ID: 3809523)(Rosa26^tdTomato^), Wt1^tm2(cre/ERT2)Wtp^/J (MGI ID: 3801682)(Wt1^CreERT2^), and Sulf1^tm1c(KOMP)Mbp^. Males Rosa26^tdTomato^; Wt1^CreERT2/+^ were bred with Rosa26^tdTomato^; Wt1^CreERT2/+^ females to generate *Wt1* knock-out embryos (homozygous Wt1^CreERT2^). Sulf1^tm1a(KOMP)Mbp^ embryonic stem cell clones (MGI ID: 5141826, MMRRC: 063141-UCD, Parental Cell Line JM8A3.N1, ES Cell Clones: DEPD00568_2_A12; DEPD00568_2_H12) were obtained from the KOMP Repository at University of California, Davis. Embryonic stem cells were injected into blastocysts, and resulting chimeras bred to achieve germline transmission of the Sulf1^tm1a(KOMP)Mbp^ allele, before crossing with flippase recombinase-expressing mice Tg(ACTFLPe)9205Dym (MGI ID: 2448985) to remove the SA/LacZ-pA/neo cassette flanked by FRT sites integrated into intron 5 of *Sulf1*. The generated Sulf1^tm1c(KOMP)Mbp^ “floxed” allele (Sulf1^fl/fl^) mice, where critical exons 6 and 7 of *Sulf1* are flanked by loxP sites, were backcrossed to a C57BL/6 background. Conditional deletion of exons 6 and 7 was expected to generate a mis-sense mutation and lead to a premature stop codon in exon 8, resulting in nonsense-mediated decay of the mutant mRNA and loss of protein expression. Sulf1^fl/fl^ mice were bred to carry Rosa26^tdTomato^; Wt1^CreERT2^ for inducible, tissue-specific lineage tracing and knock-out following tamoxifen administration (*Sulf1* iKO; deletion allele Sulf1^tm1d(KOMP)Mbp^). For embryo collection, mice were paired overnight, and females were checked the next morning for the presence of a vaginal plug. Pregnant females were administered 100 mg/kg body tamoxifen by oral gavage at E9.5, unless stated otherwise.

### Epicardial explant culture and siRNA transfection

Embryonic hearts were dissected into atrial and ventricular pieces and cultured on 1% bovine gelatine-coated chamber slides or 13mm round glass coverslips in culture medium (15% FBS, 1% Penicillin/Streptomycin, DMEM high glucose GlutaMAX). All epicardial explants were cultured at 37°C in 5% CO_2_. On day 1, explants were inspected for adherence to the coated surface and successful epicardial outgrowth. For *Wt1* KO (Wt1^CreERT2/CreERT2^) and control (Wt1^+/+^ or Wt1^CreERT2/+^) explants, fresh medium was added, and cultures were maintained for a total of 3 days. For epicardial explants from C57BL/6J wild-type hearts to be subjected to the siRNA transfection, fresh medium was provided, and siRNA and Lipofectamine RNAiMAX transfection reagents (ThermoFisher, 13778075) were prepared according to the manufacturer’s instructions. Explants were transfected with 60nM Silencer Select Negative Control (Life Technologies, 4390843) or 60nM Silencer Select Wt1 siRNA (Life Technologies, s76112) for 48 hours in culture medium (10% FBS, 1% Penicillin/Streptomycin, DMEM high glucose GlutaMAX). Finally, all explants were washed with PBS, fixed in 4% paraformaldehyde (PFA) for 15 minutes at room temperature, and stored as needed for downstream staining. Fixed explants were stored at 4°C in PBS prior to immunostaining. For RNAscope RNA fluorescence in situ hybridization (RNA FISH), fixed explants were dehydrated through a graded ethanol series (50%, 70%, and 100%) and stored at −20°C in 100% ethanol until further processing.

### Epicardial cell line culture and siRNA/pDNA transfection

MEC1, an epicardium-derived cell line ^68^ (Sigma-Aldrich, SCC187), was cultured on 0.1% bovine gelatine in MEC1 expansion medium (10% EmbryoMax ES Cell Qualified FBS ES009-M, 1% Penicillin/Streptomycin, EmbryoMAX DMEM high glucose SLM-120) at 37°C in 5% CO_2_. MEC1 cells were seeded overnight on 1% bovine gelatine-coated, assay-specific, tissue culture treated plates in MEC1 expansion medium, before being subjected to siRNA or plasmid DNA (pDNA) transfection procedures.

For siRNA transfection, cells were provided with low serum medium (2% EmbryoMax ES Cell Qualified FBS, 1% Penicillin/Streptomycin, EmbryoMAX DMEM high glucose), while siRNA and Lipofectamine RNAiMAX transfection reagents (ThermoFisher) were prepared according to the manufacturer’s instructions. MEC1 cells were transfected with 60nM Silencer Select Negative Control, 60nM Silencer Select Wt1 or 40nM Silencer Select Sulf1 siRNAs (a 1:1 combination of s109342 and s109341; Life Technologies) for 24, 48, 72 or 96 hours in low serum medium at 37°C in 5% CO_2_.

For pDNA transfection, cells were provided with serum- and antibiotic-free medium (Opti-MEM), while pDNA, P3000 enhancer and Lipofectamine 3000 transfection reagents (ThermoFisher, L3000001) were prepared as per the manufacturer’s instructions. MEC1 cells were transfected with custom vectors, 100ng pRP[En]-{Sulf1_Enh_12715347_5982_WT_SL}: miniCMV>Luciferase (VB230713-1205ffr) or 100ng pRP[En]-{Sulf1_Enh_12715347_WT1_MUT5_SL}: miniCMV>Luciferase (VB230713-1207huq) (VectorBuilder) and incubated at 37°C in 5% CO_2_ for 4 hours in serum- and antibiotic-free Opti-MEM medium (1µg pDNA/ml medium). After 4 hours, pDNA transfection medium was replaced with MEC1 expansion medium, and cells were incubated for 24 or 48 hours at 37°C in 5% CO_2_.

### Tissue collection and cryosectioning

Embryos and embryonic hearts were harvested and fixed in 4% PFA for 1.5-2 hours at room temperature. After PBS washes, tissues were equilibrated overnight at 4°C in 30% sucrose/PBS and gradually transitioned into O.C.T. embedding medium. Samples were stored at −80°C before being cryosectioned at 8-12µm thickness.

### RNAscope RNA fluorescence in situ hybridization (RNA FISH)

RNAscope Multiplex Fluorescent v2 assay (Bio-Techne, 323110) was performed on 8-12 μm cryosections or epicardial explants according to manufacturer’s instructions, with minor modifications ^2^. Briefly, for cryosections, the assay was conducted according to manufacturer’s instructions, with the following adjustments: the target retrieval step which was reduced to 10 minutes, and the Protease Plus digestion which was optimised for the embryonic stage of the tissue (i.e. E9.5-E11.5: 10 minutes; E13.5: 15 minutes; E17.5: 20 minutes) at room temperature. For epicardial explants, samples were rehydrated through a graded ethanol series (70%, 50%) into PBS, followed by digestion with protease III (1:15, in PBS) at room temperature for 10 minutes. The RNAscope assay was then performed according to manufacturer’s instructions. All catalogue probes, including the negative control, and TSA Plus fluorophores are listed below. Where required, RNA FISH was followed by immunostaining. Samples were blocked in a buffer containing 1% BSA and 10% donkey serum in PBS for 15 minutes at room temperature. Primary antibodies were added, and samples were incubated either for 1 hour at room temperature (for conjugated antibodies; cTNT, BD Biosciences, 565744, 1:100) or overnight at 4°C (for unconjugated antibodies; SMMHC, Abcam, ab125884, 1:200). After PBS washes, when required, secondary antibodies were added and incubated for 1 hour at room temperature, followed by additional PBS washes. Nuclei were stained with DAPI. Samples were mounted in Prolong Gold Antifade Mountant (ThermoFisher) with glass coverslips, then stored at 4°C until imaging. Fluorescence images of cryosections were acquired using a Leica DM6000B microscope, while confocal fluorescence images were captured with a Zeiss LSM 880 microscope. Image processing and analysis were performed using ImageJ/Fiji and CellProfiler software. Quantification was performed using custom CellProfiler pipelines, adapted from a previously published method ^145^.

|  |  |  |
| --- | --- | --- |
| RNAscope 3-plex<br>Negative Control Probe | Bio-Techne | 320871 |
| Mm-Upk3b | Bio-Techne | 568561 |
| Mm-Wt1 | Bio-Techne | 432711<br>432711-C2<br>416051-C3 |
| Mm-Sulf1 | Bio-Techne | 495411<br>495411-C2 |
| Mm-Sulf2 | Bio-Techne | 467901-C3 |
| Mm-Postn | Bio-Techne | 418581-C2 |
| Mm-Dpt | Bio-Techne | 561511 |
| Mm-Rgs5 | Bio-Techne | 430181 |
| tdTomato | Bio-Techne | 317041<br>317041-C2<br>317041-C3 |
| Mm-Pecam1 | Bio-Techne | 316721 |
| Mm-Npr3 | Bio-Techne | 502991 |
| Mm-Tcf21 | Bio-Techne | 508661-C2<br>508661-C3 |
| Mm-Fgf2 | Bio-Techne | 316851 |
| Mm-Tgfb1 | Bio-Techne | 407751-C3 |
| Mm-Tgfb2 | Bio-Techne | 406181 |
| TSA plus FITC/Cy3/Cy5<br>(1:250/ 1:1000/ 1:1500,<br>FISH) | Akoya<br>Biosciences | NEL741001KT<br>NEL752001KT |

### Immunofluorescent staining

Cryosections were rehydrated and permeabilized with 0.5% Triton X-100 in PBS for 15 minutes. Samples were blocked in staining buffer containing 4% FBS and 10% donkey serum in 0.2% Triton X-100/PBS for 1 hour at room temperature. Primary antibodies listed below were added, and samples were incubated overnight at 4°C. Following incubation, samples were washed in 0.1% Triton X-100 in PBS (PBST). Secondary antibodies were then added and incubated for 1 hour at room temperature, followed by additional PBST washes. Nuclei were stained with DAPI. Cultured epicardial explants and MEC1 cells followed the same immunofluorescence protocol, omitting the initial permeabilization step. Fluorescence images were acquired using a Leica DM6000B fluorescence microscope for cryosections, and a Molecular Devices ImageXpress Pico Automated Cell Imaging System for stained MEC1 cells in PhenoPlate™ 96-well microplates. Image processing and analysis were performed using ImageJ/Fiji, CellReporterXpress, or CellProfiler software. Quantification was completed using CellReporterXpress or a custom CellProfiler pipeline.

|  |  |  |
| --- | --- | --- |
| Anti-Wilms Tumor Protein antibody<br>[CAN-R9(IHC)-56-2] (1:100) | Abcam | ab89901 |
| Anti-CD31 antibody [2H8] (1:100) | Abcam | ab119341 |
| Anti-smooth muscle Myosin heavy<br>chain 11 antibody (1:200) | Abcam | ab125884 |
| Ki-67 Monoclonal Antibody (SolA15)<br>(1:100) | Invitrogen eBioscience | 14-5698-82 |
| p-Histone H3 Antibody (Ser 10)<br>(1:100) | Santa Cruz | sc-8656 |
| Endomucin Antibody (V.7C7) (1:100) | Santa Cruz | sc-65495 |
| Alexa Fluor® 647 Mouse Anti-Cardiac Troponin T (1:100) | BD Biosciences | 565744 |
| Anti-beta Catenin antibody [E247] - ChIP Grade (1:250) | abcam | ab32572 |
| Anti-Actin, α-Smooth Muscle - Cy3™ antibody (1:400) | Sigma-Aldrich | **C6198** |

### RNA extraction and quantitative RT-PCR

RNA was extracted from embryonic hearts using RNeasy Mini kit (Qiagen) and from cultured MEC1 cells using RNeasy Plus Micro kit (Qiagen, 74034), following the manufacturer’s instructions. RNA concentrations were measured using BMG Labtech Spectrostar Nano absorbance microplate reader equipped with an LVis Plate. cDNA was synthesized with High-Capacity cDNA Reverse Transcription kit (ThermoFisher) according to the manufacturer’s instructions and analysed by quantitative RT-PCR using the Fast SYBR Green Master Mix (ThermoFisher). Primer sequences are detailed below. The delta-delta cycle threshold (ΔΔCT) method was used for relative quantification. For embryonic hearts, ΔΔCT values were calculated relative to the mean of the housekeeping genes *Gapdh* and *Hprt1,* and referenced to E11.5 samples. For MEC1 cells, ΔΔCT values were calculated relative to the mean of the housekeeping genes *Gapdh* and *Actb,* and referenced to control siRNA samples at each time point.

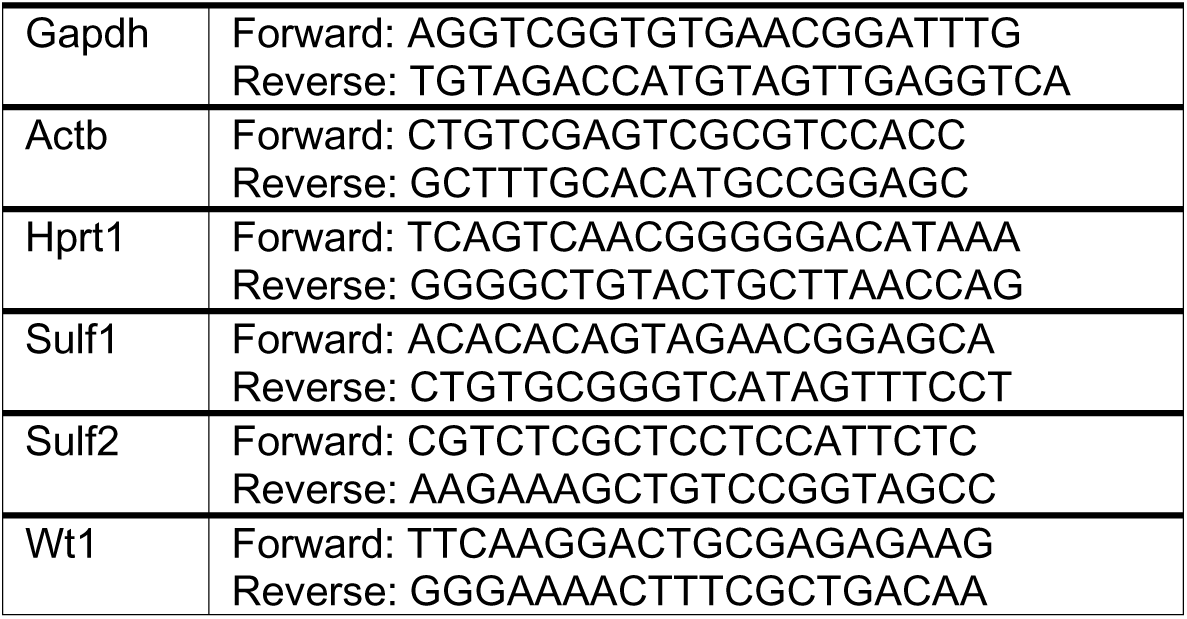

### Heart tissue dissociation and cell isolation

Whole embryonic hearts were enzymatically dissociated using the Neonatal Heart Dissociation Kit (Miltenyi Biotec, 130-098-373) and the gentleMACS Octo Dissociator with heaters (Miltenyi Biotec, 130-096-427) following the manufacturer’s instructions and as previously described ^146^. Where necessary, embryonic hearts were pooled prior to dissociation. The dissociation yielded single-cell suspensions for downstream applications, including PrimeFlow RNA Assay, FACS and 10X Genomics. The total number of cells per sample was determined using a hemocytometer and 0.4% trypan blue, with cell viability estimated to be >94%.

### PrimeFlow RNA Assay

E17.5 control hearts (Rosa26^tdTomato^; Wt1^CreERT2/+^) from littermate embryos were pooled prior to dissociation (2-3 hearts/sample) to maximise cell numbers for multiple PrimeFlow RNA staining combinations and corresponding fluorescence minus one (FMO) controls. *Sulf1* iKO hearts (Rosa26^tdTomato^; Wt1^CreERT2/+^; Sulf1^fl/fl^) were dissociated individually. Single-cell suspensions were processed using the PrimeFlow RNA Assay Kit (ThermoFisher, 88-18005-210) according to the manufacturer’s instructions, with minor modifications ^146^. Briefly, cells were surface-stained with the Zombie Aqua Fixable Viability Kit (Biolegend, 423101, 1:1000) for 30 minutes at room temperature. Samples in cell staining buffer (Biolegend, 420201) were pre-treated on ice for 5 minutes with TruStain FcX antibody to block Fcγ receptors (Biolegend, 101319, 1:50), followed by incubation with BV605 anti-CD31 antibody (Biolegend, 102427, 1:50) for 30 minutes on ice. Samples were further fixed and permeabilised using the kit reagents. Intracellular staining was performed using BV421 anti-cardiac troponin T (cTNT) antibody (BD Biosciences, 565618, 1:200) for 30 minutes on ice, followed by additional fixation, RNA target probe hybridization, and signal amplification per the manufacturer’s instructions. After signal amplification and washes, samples were stored in IC fixation buffer (1:1, cell suspension:IC fixation buffer) at 2°C-8°C, and data were acquired within 3 days. PrimeFlow catalogue probes used are detailed below. All samples were analysed using BD LSRFortessa X-20 cytometer equipped with a High Throughput Sampler (HTS), and data were processed with FlowJo software. Percent positive was calculated using Super-Enhanced Dmax Subtraction algorithm (%SED; FlowJo) for histogram comparisons. Geometric mean fluorescence intensity (gMFI) was calculated by subtracting the FMO gMFI from sample gMFI.

|  |  |  |
| --- | --- | --- |
| Wt1 | ThermoFisher Scientific | VB4-13886-PF |
| Sulf1 | ThermoFisher Scientific | VB4-3133182-PF<br>VB1-3048051-PF<br>VB6-3217745-PF |
| Sulf2 | ThermoFisher Scientific | VB1-3038415-PF<br>VB6-3208199-PF |
| Tcf21 | ThermoFisher Scientific | VB6-3197774-PF |
| Dpt | ThermoFisher Scientific | VB1-3034260-PF |
| Rgs5 | ThermoFisher Scientific | VB4-3116783-PF |
| Npr3 | ThermoFisher Scientific | VB1-3031000-PF |

### Fluorescence-activated cell sorting (FACS)

E12.5 or E13.5 hearts (Rosa26^tdTomato^; Wt1^CreERT2/+^) from littermate embryos were pooled prior to dissociation (E12.5: 7 hearts/sample; E13.5: 9-13 hearts/sample) to maximise cell numbers for FACS and corresponding FMO controls, and achieve a target sort of 50,000 epicardial cells/sample for the downstream ATAC-seq protocol. Single-cell suspensions in cell staining buffer (Biolegend) were pre-treated for 5 minutes on ice with TruStain FcX antibody, followed by surface staining with BV421 anti-CD31 (Biolegend, 102424, 1:50) and APC/Cy7 anti-Podoplanin (PDPN; Biolegend, 127418, 1:100) antibody for 30 minutes on ice, then washed with PBS before being incubated with SYTOX Green Dead Cell Stain (ThermoFisher Scientific, S34860, 1:50,000) for 5 minutes on ice. After PBS washes, cells were resuspended in sorting buffer containing 2% FBS in PBS. BD FACSAria III was used to sort 40,000-50,000 live cells negative for SYTOX Green and CD31, and positive for tdTomato and PDPN, into 0.5% BSA in PBS.

### ATAC-seq and Analysis

Live epicardial cells (CD31^−^tdTomato^+^PDPN^+^) isolated by FACS were subjected to the ATAC-seq protocol as previously described ^147^(dx.doi.org/10.17504/protocols.io.bv9mn946). In brief, 40,000-50,000 sorted cells were centrifuged at 500 ×g for 5 minutes, the supernatant was removed, and the pellet was resuspended in 50µL of ATAC Resuspension Buffer (RSB: 10mM Tris-HCl pH 7.5, 10mM NaCl, 3mM MgCl_2_) supplemented with 0.1% NP-40, 0.1% Tween-20, and 0.01% Digitonin to extract nuclei. After a 3-minute incubation, 1mL of ATAC RSB with 0.1% Tween-20 was added, mixed by inversion, and centrifuged at 500 ×g for 10 minutes. The supernatant was carefully removed, and nuclei were resuspended in 50µL of transposition mix (100mM tagmentation enzyme, Illumina, 20034197, 0.01% Digitonin, 0.1% Tween-20, 0.33X PBS, in TD buffer). The tagmentation enzyme amount was adjusted according to cell count (for 40,000 cells, the enzyme was diluted to a 0.8× concentration in TD buffer before being added to the transposition mix). Incubation and centrifugation steps were performed on ice or at 4°C, respectively, up to this point. Transposition reactions were carried out at 37°C for 30 minutes with shaking at 1,000 RPM using an Eppendorf ThermoMixer.

ATAC-seq DNA libraries were generated as previously described ^147^ (dx.doi.org/10.17504/protocols.io.bv9mn946), with minor modifications. Reactions were purified using the MinElute Reaction Cleanup Kit (Qiagen, 28204) following the manufacturer’s instructions. Pre-amplification of transposed fragments (5 cycles) was performed using NEBNext Ultra II Q5 Master Mix (NEB, M0544S) and Nextera i7 index adapters (IDT). The number of additional amplification cycles (typically 4) was determined using the qPCR method ^148^. Final PCR products were purified using a double-sided bead purification method with SPRIselect beads (0.5X volume followed by 1.3X volume of SPRI beads; Beckman Coulter, B23317). Final clean-up was performed with the MinElute Reaction Cleanup Kit (Qiagen). Libraries were assessed for quality using an Agilent TapeStation with a D1000 DNA ScreenTape assay, and quantified with the KAPA Library Quantification Kit. Next-generation sequencing (NGS) was conducted on equimolar pooled libraries using the NextSeq 500/550 High Output v2.5 kit (75 cycles) (Illumina), with paired-end reads (40 bp x 2) and 8 bp single index reads.

Sequencing data were demultiplexed using the Illumina platform and processed using Samtools, Bowtie2, Picard, Genrich, and deepTools. Nextera adaptor sequences and low-quality reads were trimmed during the bcl2fastq demultiplexing step. Reads were aligned to the mouse genome (mm10) using Bowtie2 with default parameters and a maximum fragment length of 1,000 bp. Peak calling and filtering were performed using Genrich, Samtools, and Picard. Duplicate reads were removed, and remaining reads were filtered for high quality (MAPQ ≥ 30), non-mitochondrial chromosomes, genomic blacklist regions (mm10 exclusion), and properly paired reads. Read coverage was normalized using reads per genome coverage (RPGC) and an effective genome size of 2,308,125,349. Transcription factor occupancy was predicted using TOBIAS ^80^. For visualization in the Integrative Genomics Viewer (IGV), the mean signal of two independent E13.5 samples was computed using Galaxy server’s bigwigCompare tool, with the operation set to “mean” and a bin size of 1.

### scRNA-seq and Analysis

For E17.5 single-cell RNA sequencing (scRNA-seq), a whole embryonic heart was dissociated and processed using the Chromium Single Cell 3’ Reagent Kit (v3 chemistry, 10x Genomics, PN-1000092) following the manufacturer’s instructions. Libraries were sequenced in three runs on the Illumina NextSeq 500 platform using the NextSeq® 500/550 High Output Kit v2 (150 cycles, Illumina, FC-404-200), generating 400 million reads, with a target of 8,000 cells per sample. Sequencing reads were processed with CellRanger (10x Genomics), and downstream analysis was performed in Seurat v5, as described below. Cells with more than 25% mitochondrial reads, fewer than 1,000 genes, or more than 7,000 genes were excluded from further analysis. Data were visualized using UMAP. The E10.5 scRNA-seq dataset ^48^ was processed as previously described ^2^. The E13.5 and E17.5 scRNA-seq datasets were processed as described in a prior study ^49^, with additional doublet removal performed using the scDblFinder package (based on doublet density). For batch correction and data integration of E11.5, E13.5, and E17.5 datasets, publicly available E11.5 scRNA-seq of the heart was obtained from GEO (GSE193746)^55^. Read counts from datasets (18542 cells in total) were combined and underwent shifted logarithm (log1p) normalisation. Highly variable genes were selected in a batch-aware manner using Scanpy v1.9.5 ^149^. Using the top 5000 most highly variable features, datasets were integrated using scGen v2.1.0 ^150^ with default parameters. Batch-corrected latent representation from scGen was used to calculate a new UMAP embedding. The Leiden algorithm was then used to perform cell clustering using Scanpy. The integrated epicardial cluster was identified by its expression of the markers *Upk3b*, *Upk1b*, and *Wt1*. The epicardial cluster (1093 cells) was subset and re-normalised using global scaling by the total counts per cell. Median values of gene expression were computed using Scanpy. Cell cycle phase scores (S-phase and G2M-phase) for individual cells were assigned using Seurat, and cells with *Mki67* expression levels greater than 0.5 were considered positive in the analysis. Publicly available E17.5 scRNA-seq data from endothelial cells isolated from *BmxCreERT2*;*Rosa^tdTomato^*lineage traced mouse hearts ^60^ was obtained from GEO (GSE213274) and processed as previously described ^151^.

Ligand-receptor inference analysis was performed on the E13.5 scRNA-seq dataset using CellChat ^82^. To enhance the analysis, we customized the ligand-receptor interaction database by updating the CellChatDB v1.6.0 repository with additional ligand-receptor pairs from resources such as CellPhoneDBcore (≤4.1), experimentally validated interactions from scTalkDB and STRING, and included literature-based annotations for HSPG-dependent pathways. Seurat differential expression (DE) analysis, based on the non-parametric Wilcoxon rank-sum test, was conducted on the E14.5 scRNA-seq epicardial cluster ^63^ to compare control versus Wt1^GFPCre/GFPCre^ (*Wt1* knock-out). Results were visualized in a volcano plot, with significance thresholds set at an adjusted p-value < 0.05 and fold change (FC) > 0.5.

### CUT&RUN-seq and Analysis

MEC1 cells cultured for 2 days in expansion medium were harvested from T75 flasks using 0.05% Trypsin-EDTA (2-3 minutes at 37°C). Cell viability was estimated at >95%, and cells were divided based on immunogen (1×10^6^ cells/reaction). CUT&RUN was performed using the CUT&RUN Assay Kit (Cell Signaling Technologies, 86652), following the manufacturer’s instructions with minor modifications. Cells were incubated with activated Concanavalin A magnetic beads for 10 minutes at room temperature. After washes, cell:bead suspensions were incubated overnight at 4°C with anti-WT1 (Abcam, ab89901, 1:50), anti-H3K27ac (CST, 8173, 1:100), anti-H3K4me3 (CST, 9751, 1:50), or anti-IgG (CST, 66362, 1:20) antibodies on a gentle rocker. The next day, pA/G-MNase was added, and cells were incubated at 4°C for 1 hour, followed by CaCl_2_-mediated MNase activation for DNA digestion (30 minutes at 4°C, Eppendorf ThermoMixer). Reactions were terminated with Stop buffer supplemented with 50 pg spike-in DNA per reaction (*S. cerevisiae* fragmented genomic DNA), and DNA fragments were released by incubation at 37°C for 10 minutes. DNA fragments were purified using spin columns (CST, 14209). Input DNA samples (1×10^5^ cells/reaction) were prepared using DNA extraction buffer (containing Proteinase K and RNase A) according to the manufacturer’s protocol. Samples were incubated at 55°C for 1 hour with shaking (500 RPM, Eppendorf ThermoMixer), cooled on ice, then sonicated using a Diagenode Bioruptor Pico set to “Easy Mode” (25 cycles of 30 sec on/off at 4°C). DNA was quantified using the Qubit dsDNA HS Assay Kit (Life Technologies, Q32851), and size distribution was assessed with the High Sensitivity DNA Kit (Agilent) using a Bioanalyzer. Qubit concentrations were adjusted for fragments between 50 bp and 750 bp (50 bp–1000 bp for input), calculated by normalising to the percentage of total signal in this region.

DNA libraries were prepared using the NEBNext Ultra II DNA Library Prep Kit (NEB, E7645S), with modifications guided by the protocol “Library Prep for CUT&RUN with NEBNext® Ultra™ II DNA Library Prep Kit for Illumina® (E7645) V.2” (dx.doi.org/10.17504/protocols.io.bagaibse). Less than 6 ng CUT&RUN DNA was used as input. NEBNext End Prep was performed at 20°C for 30 minutes and 50°C for 60 min. Adaptor concentrations were adjusted to input amounts (e.g., 3 µM NEBNext adaptor for 6 ng CUT&RUN DNA). Ligations were performed at 20°C for 15 min, and USER enzyme was used to cleave the uracil loop (37°C for 15 min). DNA cleanup steps used 1.75X SPRIselect beads for WT1/IgG CUT&RUN and 1.1X for histone modifications (H3K27ac, H3K4me3). To amplify the library, the ligation product was mixed with 2x Ultra II Q5 mix, universal primer and index primers (NEB, E7335S, E7500S, E7710S). PCR amplification was typically 12 - 15 cycles (WT1/IgG: 98°C for 30 sec, 12 - 15 cycles of 98°C for 10 sec, 65°C for 10 sec, final extension at 65°C for 5 min; histone modifications: 13sec annealing/extension). Double size selection (0.8X, 1.2X SPRIselect beads) was used for WT1 and IgG, while single size selection (1.1X beads) was applied for histone modifications. Input DNA samples (50 ng input) followed the standard NEBNext Ultra II DNA Library Prep Kit protocol. Libraries were assessed using a High Sensitivity DNA Kit on a on Bioanalyser instrument (Agilent) and quantified using the KAPA Library Quantification Kit (Roche, 07960336001). Next-generation sequencing (NGS) was conducted on equimolar pooled libraries using the NextSeq 500/550 High Output v2.5 kit (75 cycles) (Illumina), with paired-end reads (42 bp x 2) and 6 bp single index reads.

NGS data were demultiplexed using Illumina bcl2fastq, and processed with the nf-core/cut&run v2 pipeline using TrimGalore, Bowtie2, Samtools, Picard, deepTools, Bedtools, and SEACR. TrimGalore removed adaptor sequences, polyG+ sequences and low-quality reads, and Bowtie2 aligned reads to the mouse genome (mm10) with default parameters. Filtering was conducted with Samtools and Picard to exclude low-quality alignments (MAPQ ≥ 20 for WT1/IgG, MAPQ ≥ 10 for histone modifications), genomic blacklist regions, and duplicate reads. Peaks were called using SEACR, normalizing against IgG controls and scaling background control by a factor of 0.8. Read normalization was performed using Counts Per Million (CPM), and consensus peaks were computed from at least two biological replicates. For IGV visualization, the mean signal of two independent biological replicates was computed using Galaxy server’s bigwigCompare tool with the operation set to “mean” and bin size of 1.

Enhancer-promoter (E-P) connections were predicted using the Activity-by-Contact (ABC) model (version 0.2.2) ^81^. A custom gene annotation file was compiled to facilitate the analysis by using NCBI RefSeq annotation (build 38.1) of TSS and gene body for the mm10 genome. To infer E-P connections, 501-bp candidate enhancer elements were predicted based on MACS2-called ATAC-seq narrow peaks (using the parameters: -f BEDPE -g mm -q 0.01 --nomodel --shift −73 --extsize 146 --call-summits --cutoff-analysis --keep-dup all). Enhancer activities within the 150000 strongest elements were quantified using a combination of E13.5 epicardial ATAC-seq, H3K27ac CUT&RUN-seq of the MEC1 epicardial cell line, and TPM-normalized E13.5 epicardial gene expression (scRNA-seq). ABC scores were then computed by combining region activity and Hi-C contact frequency estimated by a power-law function of genomic distance. The power-law exponent was set to 0.87. To flag genes that are impervious to the effect of distal enhancers and exclude them from the analysis, a published list of ubiquitously expressed genes across mouse tissues ^152^ was utilised. E-P pairs were retained if their ABC scores exceeded the threshold of 0.02, corresponding to approximately 70% recall and 60% precision.

### Construction of custom vectors and pDNA purification

The consensus WT1 CUT&RUN peak sequence at the *Sulf1* locus (mm10, chr1:12,715,347-12,715,982, 636 bp, wild-type) was cloned into a mammalian enhancer-testing vector containing a miniCMV promoter and luciferase reporter gene (WT_SL; VectorBuilder Inc.). For loss of functional TF binding site analysis, a mutant sequence (WT1_MUT5_SL) was created, in which the two predicted WT1 binding motifs (JASPAR Wt1_MA1627.1) were mutated by altering five nucleotides each. To verify these mutations and assess their impact, *in silico* motif scanning was performed using MEME Suite tools (MAST for motif scanning, Tomtom for motif comparison), ensuring that WT1 binding was abolished without creating new significant binding sites for other transcription factors expressed in epicardial cells.

Custom vectors, pRP[En]-{Sulf1_Enh_12715347_5982_WT_SL}: miniCMV>Luciferase (VB230713-1205ffr) and pRP[En]-{Sulf1_Enh_12715347_WT1_MUT5_SL}: miniCMV>Luciferase (VB230713-1207huq), were generated by VectorBuilder and supplied as E. coli stocks. Bacteria harbouring plasmid DNA vectors were plated on LB-agar containing ampicillin and incubated overnight at 37°C. Individual colonies were picked and expanded in culture, and plasmid DNA was extracted using the PureLink™ HiPure Plasmid Maxiprep Kit (Invitrogen). The incorporation of the enhancer insert was confirmed by restriction digest using NheI-HF, Bsu36I, and XhoI (NEB).

### Heparan sulfate (HS) staining and analysis

Following siRNA transfection of MEC1 cells (48 hours) in PhenoPlate 96-well, flat, optically clear-bottom, black TC-treated microplates (Revvity), the medium was removed, and cells were washed with PBS and fixed with 4% PFA for 10 minutes at room temperature. Cells were then washed and stored in PBS at 4°C prior to HS staining targeting 2-*O*, *N*- and 6-*O*-sulfated epitopes. Cells were blocked in staining buffer containing 0.1% (w/v) BSA in PBS for 30min at room temperature. VSV-tagged phage-display single-chain variable fragment (ScFv) antibodies, AO4B08 (^71^, 1:25) or HS3B8 (^72^, 1:25) were added, and samples were incubated for 1 hour at room temperature. After washing with PBS, samples were incubated with a Cy3-conjugated anti-VSV glycoprotein antibody (Sigma-Aldrich, C7706, 1:100) for 1 hour at room temperature, followed by additional PBS washes. Stained cells were then rinsed in 100% Ethanol, then washed again in PBS, before nuclei were stained with DAPI. Fluorescence images were captured using the Molecular Devices ImageXpress Pico Automated Cell Imaging System. Each condition was tested in single wells, with five images per well captured using a 20X objective. Image analysis was conducted using ImageJ/Fiji and CellProfiler software. Quantification was performed with a custom CellProfiler pipeline. Briefly, nuclei segmentation was achieved through thresholding on DAPI staining intensities. HS staining was detected by thresholding on fluorescence intensity within areas co-localising with cellular regions. For each image, both the total HS-positive area (e.g., AO4B08-positive area) and total intensity were measured. Corrected total cellular fluorescence (CTCF) was calculated as: CTCF = Total Intensity – (Area occupied by cells x Mean Background Intensity). This value was normalized by nuclei count to account for cell density differences, and images containing fewer than 700 nuclei were excluded from analysis.

### Luciferase Activity Assay

MEC1 cells were seeded into 96-well, flat, clear-bottom, white TC-treated microplates and transfected with siRNAs for 24 hours, followed by plasmid DNA (pDNA) transfection (4 hours) and an additional 24 or 48 hours at 37°C in 5% CO_2_. Luciferase activity was measured using the Bright-Glo luciferase assay system (Promega, E2610) following the manufacturer’s protocol. Briefly, the 96-well plates were removed from the incubator and allowed to equilibrate at room temperature for 5 minutes. Bright-Glo reagent was then added to each well in a 1:1 ratio with the medium (100 µL reagent:100 µL medium). After a 2-minute incubation at room temperature, luminescence was measured using a BMG Labtech CLARIOstar luminometer microplate reader.

### Proliferation Assay

Following siRNA transfection of MEC1 cells (48 hours) in PhenoPlate 96-well, flat, optically clear-bottom, black TC-treated microplates (Revvity), the medium was removed, and cells were washed with DMEM. The medium was replaced with EmbryoMax DMEM containing 2% KnockOut Serum Replacement and 1% Penicillin/Streptomycin. Cells were incubated at 37°C in 5% CO₂ for 4-5 hours before adding recombinant proteins at the following concentrations: 10 ng/mL hTGF-β1, 10 ng/mL hTGF-β2, 20 ng/mL (50 ng/mL) hFGF2, 50 ng/mL hPDGF-BB, 100 ng/mL (10 ng/mL) hMidkine, 100 ng/mL (20 ng/mL) hPleiotrophin, 100 ng/mL (500 ng/mL) h/mWnt-5a, 50 ng/mL (100 ng/mL) mVEGF-B, 50 ng/mL hIGF-I, 20 ng/mL hR-Spondin-3, 50 ng/mL (10 ng/mL) hBMP-10, 100 ng/mL (50 ng/mL) hIGF-BP4, and 50 ng/mL h/m/rBMP-2 Additional concentrations tested in brackets. An hour before the 24-hour treatment at 37°C in 5% CO_2_ ended, 10 µM EdU (ThermoFisher) was added to the medium, and cells were incubated for 1 hour at 37°C in 5% CO_2_. After incubation, cells were washed with PBS, fixed with 4% paraformaldehyde (PFA) for 15 minutes at room temperature, and stored in PBS at 4°C after additional washes. For EdU detection, cells were permeabilized with 0.5% Triton-X-100/PBS for 20 minutes at room temperature, then stained using the Click-iT EdU Alexa Fluor 488 Imaging Kit (5 µM Alexa Fluor azide, ThermoFisher Scientific, C10337), following the manufacturer’s protocol. Fluorescence images were captured using the Molecular Devices ImageXpress Pico Automated Cell Imaging System. Each condition was tested in duplicate wells, with one image per well captured using a 4X objective. Image analysis was conducted using ImageJ/Fiji, CellReporterXpress, or CellProfiler software. DAPI staining was used to identify nuclei, and a threshold was set to detect EdU-positive nuclei, quantifying cell proliferation. Quantification was performed with CellReporterXpress or a custom CellProfiler pipeline. Images containing fewer than 100 cells were excluded from analysis.

|  |  |  |
| --- | --- | --- |
| Recombinant Human TGF-β1 (CHO derived) | PeproTech | Cat# 100-21C |
| Recombinant Human TGF-β2 (HEK293 derived) | PeproTech | Cat# 100-35B |
| Recombinant Human FGF-basic (146 a.a.) | PeproTech | Cat# 100-18C |
| Recombinant Human PDGF-BB | PeproTech | Cat# 100-14B |
| Recombinant Human Midkine | PeproTech | Cat# 450-16 |
| Recombinant Human Pleiotrophin | PeproTech | Cat# 450-15 |
| Recombinant Human/Mouse Wnt-5a | Bio-Techne | Cat# 645-WN-010 |
| Recombinant Mouse VEGF-B 167 | Bio-Techne | Cat# 2595-VE-025 |
| Recombinant Human IGF-I | PeproTech | Cat# 100-11 |
| Recombinant Human R-Spondin-3 | PeproTech | Cat# 120-44 |
| Recombinant Human BMP-10 | PeproTech | Cat# 120-40 |
| Recombinant Human IGF-BP4 | PeproTech | Cat# 350-05B |
| Recombinant Human/Murine/Rat BMP-2 (CHO derived) | PeproTech | Cat# 120-02C |

### Epithelial-Mesenchymal Transition Assay

Following siRNA transfection of MEC1 cells (48 hours) in PhenoPlate 96-well, flat, optically clear-bottom, black TC-treated microplates (Revvity), the medium was removed, and cells were washed with DMEM. The medium was replaced with EmbryoMax DMEM containing 2% KnockOut Serum Replacement and 1% Penicillin/Streptomycin. Recombinant proteins were added at the following concentrations: 10 ng/mL hTGF-β1, 10 ng/mL hTGF-β2, 20 ng/mL (50 ng/mL) hFGF2, 50 ng/mL hPDGF-BB, 100 ng/mL (10 ng/mL) hMidkine, 100 ng/mL (20 ng/mL) hPleiotrophin, 100 ng/mL (500 ng/mL) h/mWnt-5a, 50 ng/mL (100 ng/mL) mVEGF-B, 50 ng/mL hIGF-I, 20 ng/mL hR-Spondin-3, 50 ng/mL (10 ng/mL) hBMP-10, 100 ng/mL (50 ng/mL) hIGF-BP4, and 50 ng/mL h/m/rBMP-2 Additional concentrations tested in brackets. After a 24-hour incubation at 37°C in 5% CO_2_, cells were washed with PBS and fixed with 4% paraformaldehyde (PFA) for 10 minutes at room temperature. Cells were then washed and stored in PBS at 4°C prior to immunostaining, conducted as outlined in this study. Fluorescence images were captured using the Molecular Devices ImageXpress Pico Automated Cell Imaging System. Each condition was tested in duplicate wells, with five images per well captured using a 20X objective. Image analysis was conducted using ImageJ/Fiji or CellProfiler software. Quantification was performed with a custom CellProfiler pipeline. Briefly, the pipeline classified cells based on signal localisation, structure, texture and fluorescence intensity of β-CATENIN and α-SMA staining, categorising cells into four populations: 1. positive for membrane β-CATENIN and negative for α-SMA filaments (β-cat^hi^ α-SMA^−^), 2. disrupted membrane β-CATENIN and negative for α-SMA filaments (β-cat^dis^ α-SMA^−^), 3. negative for membrane β-CATENIN and low or negative for α-SMA filaments (β-cat^−^ α-SMA^−/lo^), d) disrupted/absent membrane β-CATENIN and 4. positive for α-SMA filaments (β-cat^−/dis^ α-SMA^hi^). Nuclei and cell segmentation was achieved through thresholding on DAPI and β-CATENIN staining intensities, respectively. Cell body boundaries were estimated using β-CATENIN staining and autofluorescence, and outlined by propagating from the segmented nuclei. The cytoplasm was defined by subtracting the nuclear area from the cell body area. Tubeness and line structure filters were applied to enhance and segment α-SMA filaments. Circularity was measured to identify and exclude artifacts, such as fluorescent antibody precipitates. Additional thresholds were applied to refine classification based on: 1. mean intensity of a-SMA filaments, 2. mean intensity of β-CATENIN at cell boundaries, and 3. average sum texture of β-CATENIN at cell boundaries. Images containing fewer than 100 cells were excluded from analysis.

### Western Blot and analysis

Following siRNA transfection of MEC1 cells (48 hours) in 12-well plates, the medium was removed, and cells were washed with DMEM. The medium was replaced with EmbryoMax DMEM containing 2% KnockOut Serum Replacement and 1% Penicillin/Streptomycin. Cells were incubated at 37°C in 5% CO₂ for 4 - 5 hours prior to adding recombinant hTGF-β1 protein at a concentration of 10 ng/mL. Cells were incubated with hTGF-β1 for various time points (15, 30, 60, 90, 120, 150 minutes) at 37°C in 5% CO_2_. Following incubation, cells were washed with cold PBS and lysed using RIPA buffer supplemented with 5mM NEM, 1mM DTT, 0.125U/mL Benzonase, and protease/phosphatase inhibitor cocktails (Sigma, Roche). Protein concentrations were quantified using the Pierce BCA Protein Assay Kit - Reducing Agent Compatible (Thermo Scientific) and measured using BMG Labtech Spectrostar Nano absorbance microplate reader. Samples were reduced, denatured, and loaded onto NuPAGE 10% Bis-Tris Protein Gels (ThermoFisher Scientific). After SDS-PAGE, proteins were transferred to PVDF membranes. Blots were blocked in 8% skimmed milk/TBST (TBS + 0.1% Tween 20, Sigma) for 1 hour, washed with TBST, and incubated overnight at 4°C with primary antibodies, anti-pSmad2 (CST, 3108,1:1000), anti-Smad2 (CST, 3103,1:1000) and anti-β-Actin (Abcam, ab8224, 1:2500) diluted in 5% BSA/TBST. The following day, blots were washed and incubated with HRP-conjugated secondary antibodies (diluted 1:2500-1:5000 in 5% skimmed milk/TBST) for 1-2 hours at room temperature. Blots were washed in TBST and developed using Immobilon Crescendo Western HRP substrate, then imaged on a ChemiDoc system. Blots were stripped and re-probed to detect different protein targets. Western blot band intensities were quantified using ImageJ software.

### Statistical Analysis

Statistical analyses were performed in GraphPad Prism v10 software. An unpaired t-test with Welch’s correction, when necessary, was applied for comparisons between two groups. For comparisons involving more than two groups, one-way ANOVA with Dunnett’s or Šídák’s multiple comparison test, when necessary, was used. One-Way ANOVA and Fisher’s LSD post-hoc test was used when multiple comparison correction was not applicable as questions were set for specific groups. Two-Way ANOVA with Dunnett’s multiple comparison test was employed for comparisons involving more than two groups with two independent variables. Results are reported as mean ± s.e.m. Significance was defined as *P* < 0.05. Statistical tests, biological/independent replicates, and *P* values are provided in the figure legends.

## Data availability

scRNAseq (GSE299996), ATAC-seq (GSE299995) and CUT&RUN-seq (GSE299998) data were deposited in the Gene Expression Omnibus (GEO). This paper analyses existing, publicly available data, accessible at GSE76118, GSE205797, GSE193746 and GSE213274. Correspondence and requests for materials should be addressed to A.N.R and N.S.

## Code availability

Scripts and custom pipelines are available on GitHub at https://github.com/RedpathAN.

## Acknowledgements

We thank Dr Madeleine Lemieux (Bioinfo) for processing 10X data with Cellranger; Dr Neil Ashley, WIMM Single cell facility, for 10X libraries and sequencing, the Dunn School and Jenner Institute Flow Cytometry facility for advice and assistance, and Biomedical Services staff for animal husbandry. We thank Dr. Toin van Kuppevelt (Amsterdam UMC, The Netherlands) for kindly providing anti-HS phage display antibodies, AO4B08 and HS3B8. We thank Prof. Paul Riley (IDRM, University of Oxford, UK) for helpful comments that improved the clarity of the paper. This work was funded by the British Heart Foundation (BHF) Project grant (N.S.: PG/16/27/32114); BHF Ian Fleming Fellowship (N.S.: FS/19/32/34376); BHF Intermediate Basic Science Research Fellowship (J.M.V.: FS/19/31/34158); Oxford BHF Centre of Research Excellence (J.M.V., A.N.R. and I-E.L.: RE/18/3/34214); BHF Centre of Regenerative Medicine, Oxford (RM/13/3/30159); BHF Immediate Research Fellowship (I.R.M.: FS/IPBSRF/23/27085)

## Author Contributions

Conceptualisation, A.N.R. and N.S.; Methodology, A.N.R, I-E.L., J.M.V.; Investigation, A.N.R., I-E.L., L.H., Q.D., I.R.M., T.C., J.M.V.; Resources, T.v.K.; Formal Analysis, A.N.R.; Visualisation, A.N.R.; Writing – Original Draft, A.N.R.; Writing – Review and Editing, A.N.R., I-E.L., Q.D., I.R.M., J.M.V., N.S.; Funding Acquisition, A.N.R., I-E.L., J.M.V., N.S.; Supervision, A.N.R. and N.S.

## Additional information

Extended data is available for this paper.

