## Extended Data Figures for "Essential regulation of heparan sulfate proteoglycan signalling controls cell behaviour to support cardiac development"

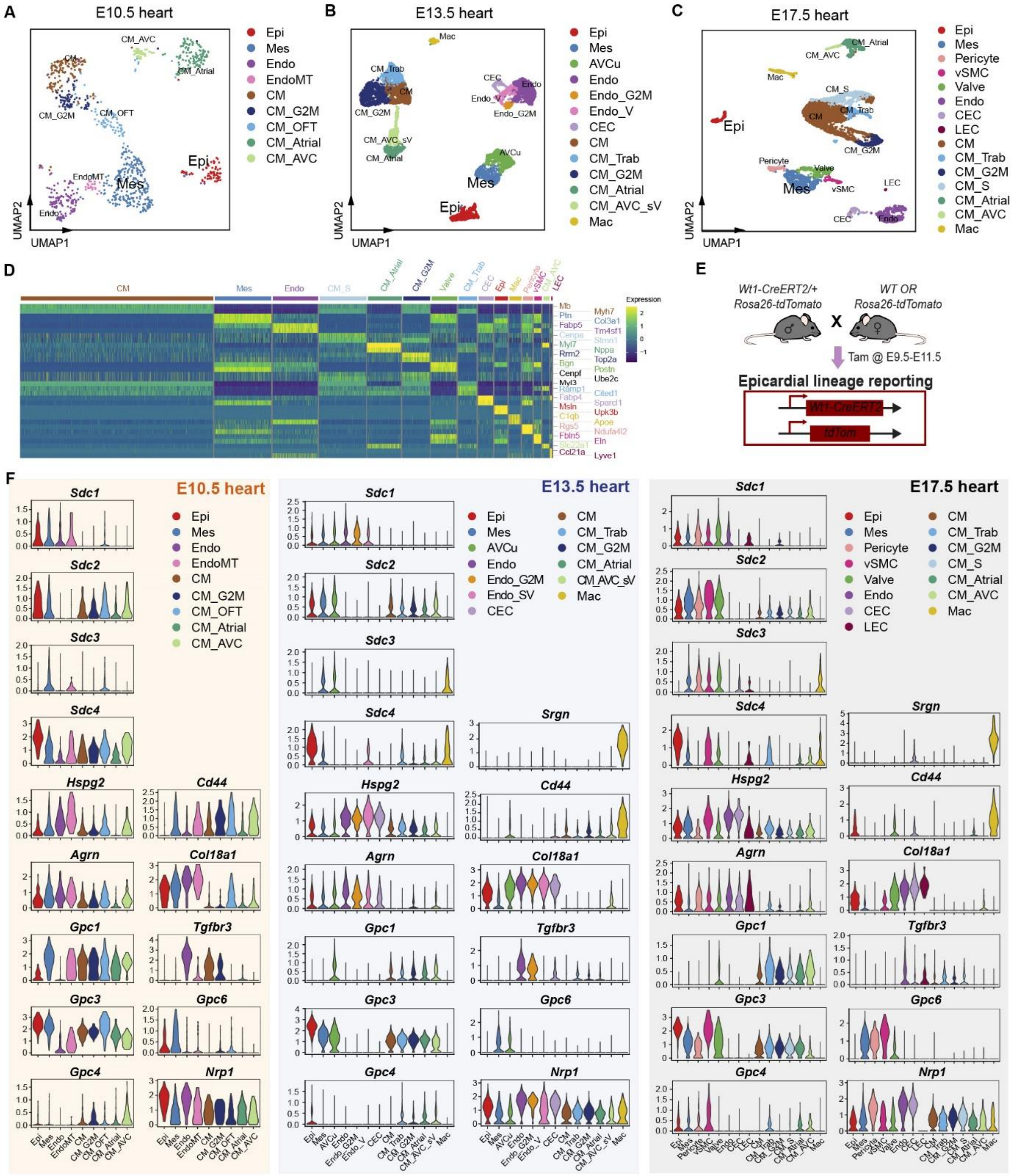

**Extended Data Fig.1 Transcriptional profiling of heparan sulfate proteoglycans in the developing mouse heart**

(A) Uniform manifold approximation and projection (UMAP) plot showing the major clusters in E10.5 hearts (total 928 cells, n = 2 batches). (B) UMAP plot showing the major clusters in E13.5 hearts (total of 6,939 cells, n = 3 hearts). (C) UMAP plot showing the major clusters in a E17.5 heart (total of 6,341 cells, n = 1 heart). (D) Heatmap of the top 2 differentially expressed genes for each cluster in the E17.5 scRNA-seq data. High expression is indicated in yellow. (E) Generation of epicardial-lineage reporting hearts by crossing heterozygous carrier of Wt1-CreERT2 allele and Rosa26-tdTomato allele. Pregnant females induced with tamoxifen at E9.5 - E11.5 to label the newly-formed epicardium (E10.5 - E11.5) and trace its derivatives. (F) Violin plots showing relative expression of ‘full-time’ and ‘part-time’ heparan sulfate proteoglycans (HSPG) in the developing mouse embryo at E10.5, E13.5 and E17.5.

*AVCu, atrioventricular cushion; CEC, coronary endothelial cells; CM, ventricular cardiomyocytes; CM_Atrial, atrial cardiomyocytes; CM_AVC, atrioventricular canal cardiomyocytes; CM_AVC_sV, atrioventricular canal and sinus venosus cardiomyocytes; CM_OFT, outflow tract cardiomyocytes; CM_S, S-G2M phase progressing cardiomyocytes; CM_Trab, trabecular cardiomyocytes; Endo, endocardial cells; Endo_V, valvular endothelial and endocardial cells; EndoMT, endocardial-to-mesenchymal transition; Epi, epicardium; G2M, proliferating; LEC, lymphatic endothelial cells; Mac, macrophages; Mes, mesenchyme; Valve, valve mesenchyme; vSMC, vascular smooth muscle cells*.

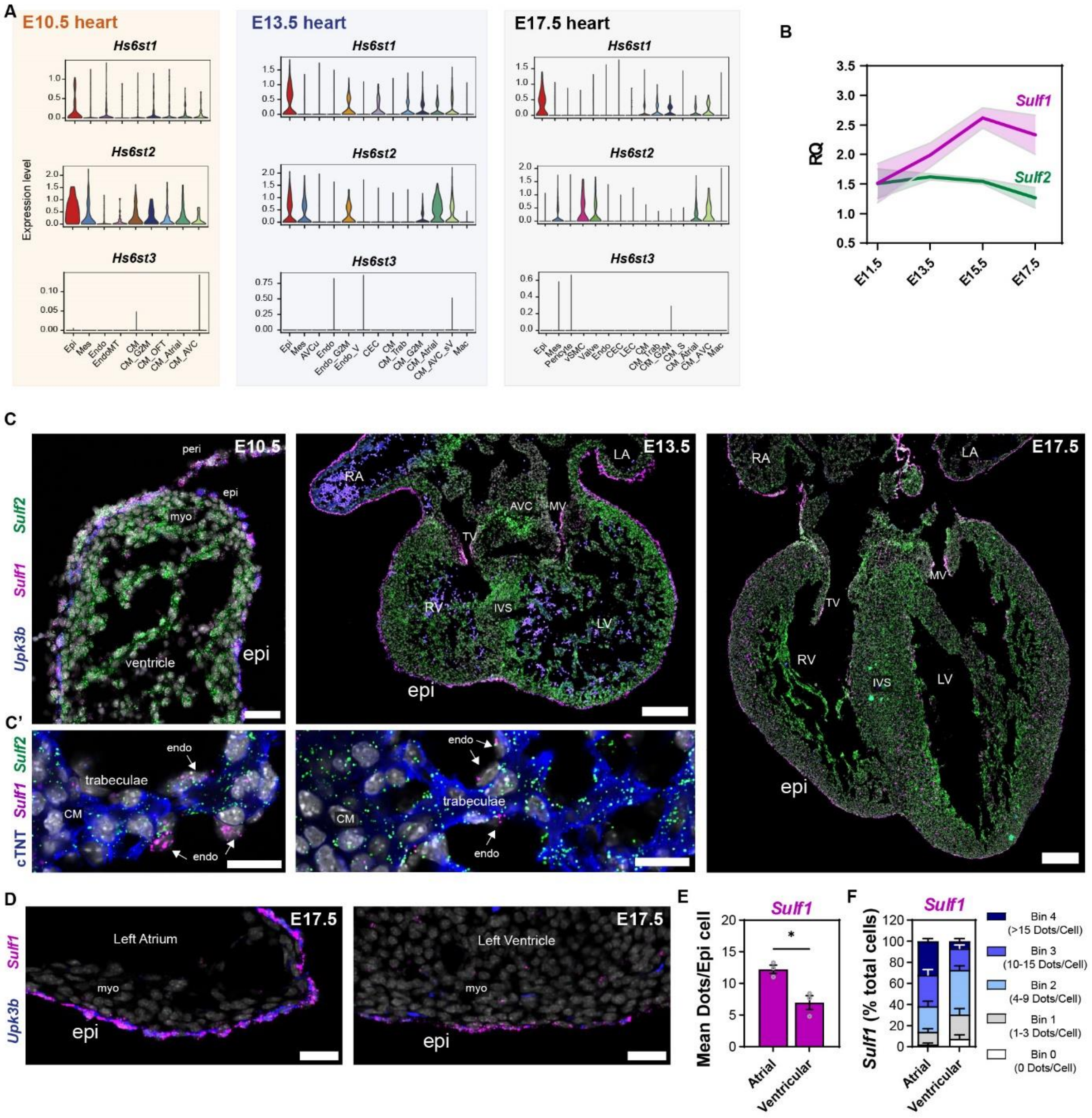


**Extended Data Fig. 2 Transcriptional profiling of heparan sulfate 6-*O*-sulfotransferases and 6-*O*-endosulfatases in the developing mouse heart**

(A) Violin plots showing relative expression of *Hs6st1*, *Hs6st2* and *Hs6st3* in the developing mouse embryonic heart at E10.5, E13.5 and E17.5. (B) qRT-PCR analysis of *Sulf1* and *Sulf2* transcript expression in whole heart lysates at E11.5, E13.5, E15.5 and E17.5. Values normalized to *Hprt* and *Gapdh*, data are represented as mean ± SEM with shaded error bars (n = 4 hearts). (C) Fluorescence ISH on cryosections from E10.5 embryos, E13.5 and E17.5 hearts for *Upk3b*, *Sulf1* and *Sulf2*, showing expression of *Sulf1* in the pericardium, epicardium and derivatives, while widespread expression of *Sulf2* is detected in the pericardium and myocardium. *Sulf1* is also detected in the forming atrioventricular valves, and early endocardium. Insets show cardiomyocytes expressing *Sulf2*, and endocardial cells expressing *Sulf1* in the trabeculated myocardium. Scale bars: 50 µm (E10.5), 200 µm (E13.5, E17.5) and inset 20 µm. Images representative of n = 7 embryos for E10.5, n = 4 hearts for

E13.5, and n = 5 hearts for E17.5. (D) Fluorescence ISH of E17.5 heart cryosections for *Upk3b* and *Sulf1*, and corresponding quantification (E) of *Sulf1* in atrial and ventricular epicardial cells. Scale bars: 20 μm. Images representative of n = 3 hearts. Data are represented as mean ± SEM (n = 3 hearts). Unpaired t-test with Welch's correction (*) P <0.05. (F) Percentage of epicardial cells expressing *Sulf1*, grouped into 5 bins based on the number of dots (individual RNA molecules) per cell. Data are represented as mean ± SEM (n = 3 hearts).

*AVCu, atrioventricular cushion; CEC, coronary endothelial cells; CM, ventricular cardiomyocytes; CM_Atrial, atrial cardiomyocytes; CM_AVC, atrioventricular canal cardiomyocytes; CM_AVC_sV, atrioventricular canal and sinus venosus cardiomyocytes; CM_OFT, outflow tract cardiomyocytes; CM_S, S-G2M phase progressing cardiomyocytes; CM_Trab, trabecular cardiomyocytes; Endo, endocardial cells; Endo_V, valvular endothelial and endocardial cells; EndoMT, endocardial-to-mesenchymal transition; Epi, epicardium; G2M, proliferating; LEC, lymphatic endothelial cells; Mac, macrophages; Mes, mesenchyme; Valve, valve mesenchyme; vSMC, vascular smooth muscle cells, peri, pericardium; epi, epicardium; myo, myocardium; RA, right atrium; RV, right ventricle; IVS, interventricular septum; LA, left atrium; LV, left ventricle; AVC, atrioventricular canal; TV, tricuspid valves; MV, mitral valves.*

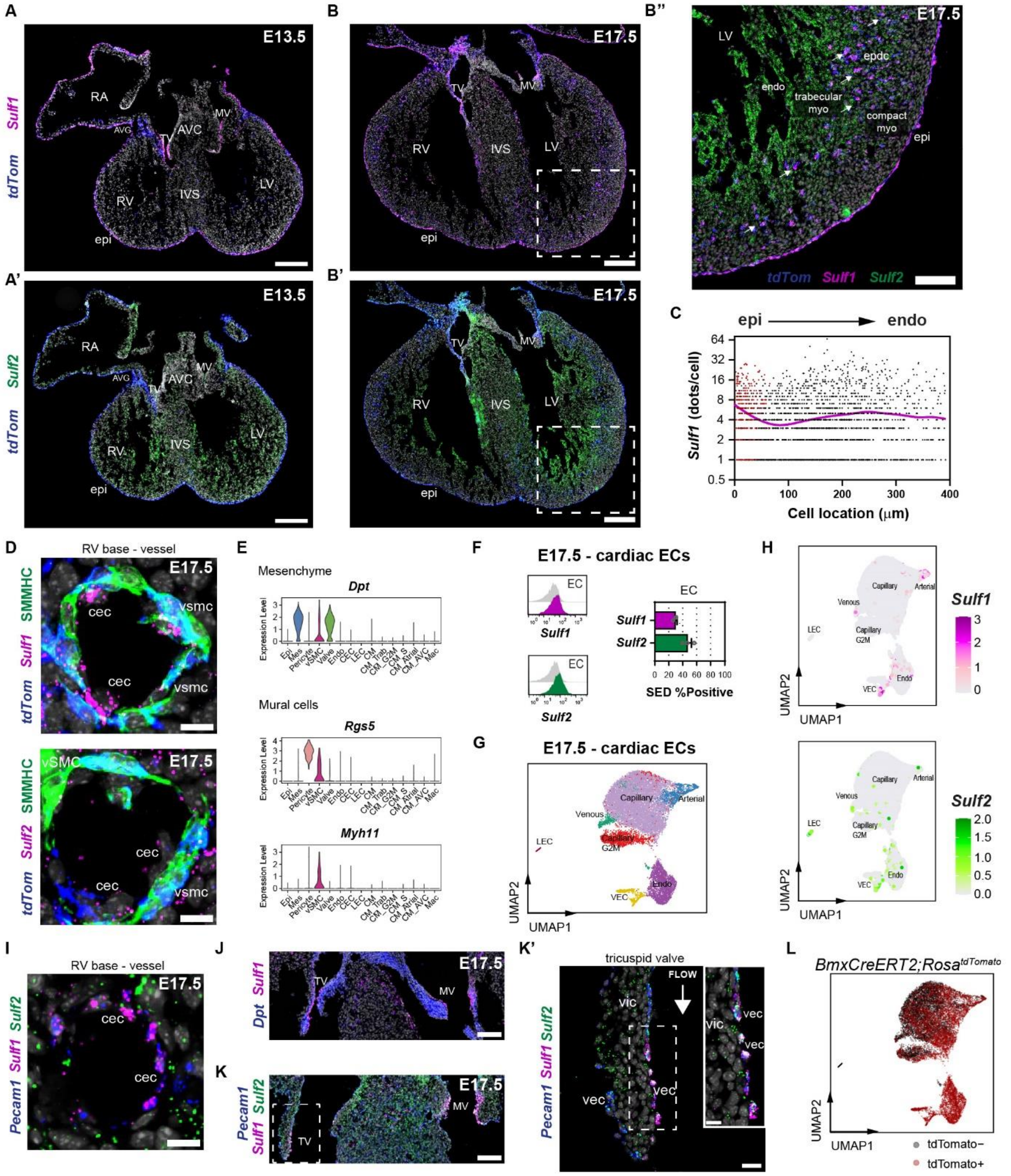


**Extended Data Fig. 3 6-*O*-endosulfatases are distinctively expressed in cardiac mesenchyme and endothelial cell compartments**

(A) Fluorescence ISH of E13.5 heart cryosections for *tdTomato*, *Sulf1* and *Sulf2*, showing majority expression of *Sulf1* in the epicardium, and atrioventricular valves. Scale bars: 250 µm. Images representative of n = 5 hearts. (B) Fluorescence ISH of E17.5 heart cryosections for *tdTomato*, *Sulf1* and *Sulf2* showing expression of *Sulf1* and *Sulf2* in epicardium-derived mesenchymal cells invading the myocardium. (B'') *Sulf1* demonstrates a trend, from high expression in the epicardium, to low expression in EPDCs located in the compact myocardium, and high expression in EPDCs located proximal to the trabecular myocardium. *Sulf2* demonstrates a gradient, from high expression in the trabecular myocardium to low expression in the compact myocardium. Scale bars: 250 µm and inset 100 µm. Images representative of n = 5 hearts. (C) Scatter plot showing quantification of *Sulf1* expression in ventricular epicardial and derivative *tdTomato* positive cells (using Fluorescence ISH images; ROI apex and mid). Data of individual cells from n = 4 hearts, and two ROI images per heart. Cell location is indicated by the Y center coordinate within each ROI image: values of $y$<100µm predominantly correspond to cells in the epicardium and subepicardium, while higher $y$ values reflect increasing depth into the compact and trabecular myocardium. A magenta LOWESS smoothing line models *Sulf1* expression in epicardium-lineage cells along their spatial distribution from the epicardium toward the endocardium. Epicardial cells are shown in red. (D) Fluorescence ISH of E17.5 heart cryosections for *tdTomato*, *Sulf1*, *Sulf2* and SMMHC, showing low expression of *Sulf1* and *Sulf2* in epicardium-derived smooth muscle cells. Scale bars: 10 µm. Images representative of n = 2 hearts. (E) Violin plots showing relative expression of markers used to distinguish mesenchyme and mural cells in E17.5 hearts. (F) Flow cytometry analysis of E17.5 endothelial cells in the heart, and corresponding quantification of percentage positive for *Sulf1* and *Sulf2*. Endothelial cells were selected by gating $CD31^+$ cells. Fluorescence minus one (FMO) control shown (grey). Data are represented as mean ± SEM (pooled 2 hearts per replicate; n = 3). (G) UMAP plot showing the major clusters in E17.5 endothelial cells from *BmxCreERT2*;*Rosa*$^{tdTomato}$ hearts. (H) UMAP feature plot showing expression of *Sulf1* and *Sulf2*, in individual endothelial cells of E17.5 *BmxCreERT2*;*Rosa*$^{tdTomato}$ hearts. (I) Fluorescence ISH of E17.5 heart cryosections for *Sulf1*, *Sulf2* and *Pecam1*, showing expression of *Sulf1* enriched in coronary endothelial cells of macrovessels. Scale bars: 10 µm. Images representative of n = 3 hearts. (J-K) Fluorescence ISH of E17.5 heart cryosections for *Sulf1, Dpt*, *Sulf2* and *Pecam1*, showing co-expression of *Sulf1 and Sulf2* in atrioventricular valve endothelial cells. (K') White arrow represents blood flow. Scale bars: 20 µm and inset 10 µm. Images representative of n = 3 hearts. (L) UMAP plot showing *tdTomato* positive and negative endothelial cells in the E17.5 *BmxCreERT2*;*Rosa*$^{tdTomato}$ hearts. *tdTomato* positive ECs mark the endocardial lineage achieved through temporal labelling (Tamoxifen induction at E9.5).

*CEC, coronary endothelial cells; CM, ventricular cardiomyocytes; CM_Atrial, atrial cardiomyocytes; CM_AVC, atrioventricular canal cardiomyocytes; CM_S, S-G2M phase progressing cardiomyocytes; CM_Trab, trabecular cardiomyocytes; Endo, endocardial cells; Epi, epicardium; G2M, proliferating; LEC, lymphatic endothelial cells; Mac, macrophages; Mes, mesenchyme; Valve, valve mesenchyme; vSMC, vascular smooth muscle cells; epi, epicardium; myo, myocardium; RA, right atrium; RV, right ventricle; IVS, interventricular septum; LV, left ventricle; AVC, atrioventricular canal; AVG, atrioventricular groove; TV, tricuspid valves; MV, mitral valves; epdc, epicardium-derived cells; vec, valve endothelial cells; vic, valve interstitial cells.*

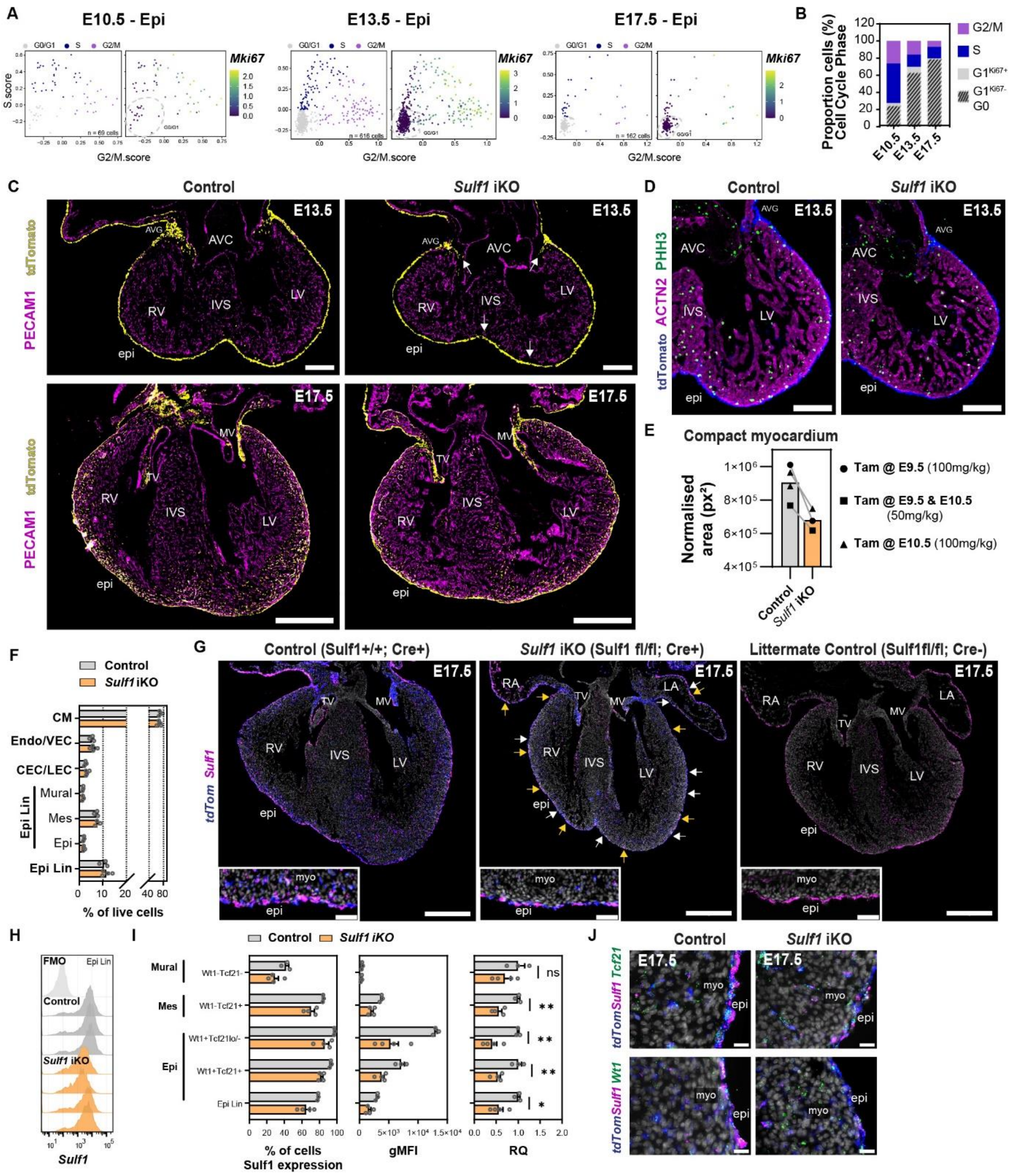


**Extended Data Fig. 4** ***Sulf1*** **iKO hearts demonstrate mosaicism and compensation as development progresses**

(A) Scatter plot showing the distribution of epicardial cells in the indicated cell cycle phases at E10.5, E13.5 and E17.5. Feature plot representing range of expression of *Mki67*, further separating quiescent and slowly-dividing epicardial cells. (B) Percentage of epicardial cells in the indicated cell cycle phases. (C) Immunofluorescence images showing tdTomato-labelled epicardium and derivatives in control and *Sulf1* iKO embryonic mouse hearts at E13.5 and E17.5. White arrows highlight reduced invasion of EPDCs at the AVG, subepicardium and IVS. Scale bar, 200 µm (E13.5) and 500 µm (E17.5). Images representative of n = 2 hearts. (D) Immunofluorescence images showing mitotic cells expressing PHH3 in control and *Sulf1* iKO embryonic mouse hearts at E13.5. Scale bar, 200µm. Images representative of n = 2 hearts. (E) Quantification of the compact myocardial area in the left ventricle of control and *Sulf1* iKO hearts at E13.5. Compact myocardial area was corrected using a normalization factor based on the length of the left ventricular surface. Data from littermate pairs are connected by lines, and different symbols indicate the tamoxifen regimens used. (F) Flow cytometry analysis of control and *Sulf1* iKO hearts at E17.5, and corresponding quantification of epicardial-lineage cells and other major cardiac cell types. The epicardial lineage was selected by gating $CD31^-cTnT^-tdTomato^+$, with subsequent gating of mesenchymal cells ($Dpt^+Rgs5^-$) and mural cells ($Rgs5^+$). Endothelial cells were selected by gating $CD31^+Npr3^{-/low}$ for coronary (CEC) and lymphatic endothelial cells (LEC), and $CD31^+Npr3^+$ for endocardial (Endo) and valve endothelial cells (VEC). Cardiomyocytes were selected by gating cells positive for cTNT. Data are represented as mean ± SEM (pooled 2 hearts per control replicate; n = 3-4). Multiple Unpaired t-test with Welch's correction; no significance. (G) Fluorescence ISH of E17.5 control and *Sulf1* iKO heart cryosections for *tdTomato* and *Sulf1*, showing a mosaic expression profile of *Sulf1* in the epicardium. White arrows highlight epicardial cells with undetectable *Sulf1* transcripts, and yellow arrows highlight epicardial cells expressing *Sulf1* comparable to control. Scale bars: 500 µm and inset 50 µm. n = 1. (H) Flow cytometry analysis of control and *Sulf1* iKO hearts at E17.5, and corresponding expression of *Sulf1* in epicardial-lineage cells (Epi Lin). Fluorescence minus one (FMO) control shown (light grey). (I) Flow cytometry analysis of control and *Sulf1* iKO hearts at E17.5, and corresponding quantification of percentage positive epicardial-lineage cells for *Sulf1*. Population expression of *Sulf1* quantified as geometric mean fluorescence intensity (gMFI), and expressed relative to control hearts (RQ). Data are represented as mean ± SEM (pooled 2 hearts per control replicate; n = 3-4). Multiple Unpaired t-test with Welch's correction (*) P <0.05, (**) P <0.01 and ns, not significant. (J) Fluorescence ISH of E17.5 control and *Sulf1* iKO heart cryosections for *tdTomato*, *Sulf1*, *Tcf21* and *Wt1*, showing expression of *Sulf1* in the epicardium and its derivative mesenchymal cells invading the myocardium. Scale bars: 20 µm. n = 1.

*epi, epicardium; myo, myocardium; RA, right atrium; RV, right ventricle; IVS, interventricular septum; LV, left ventricle; AVC, atrioventricular canal; AVG, atrioventricular groove; TV, tricuspid valves; MV, mitral valve.*

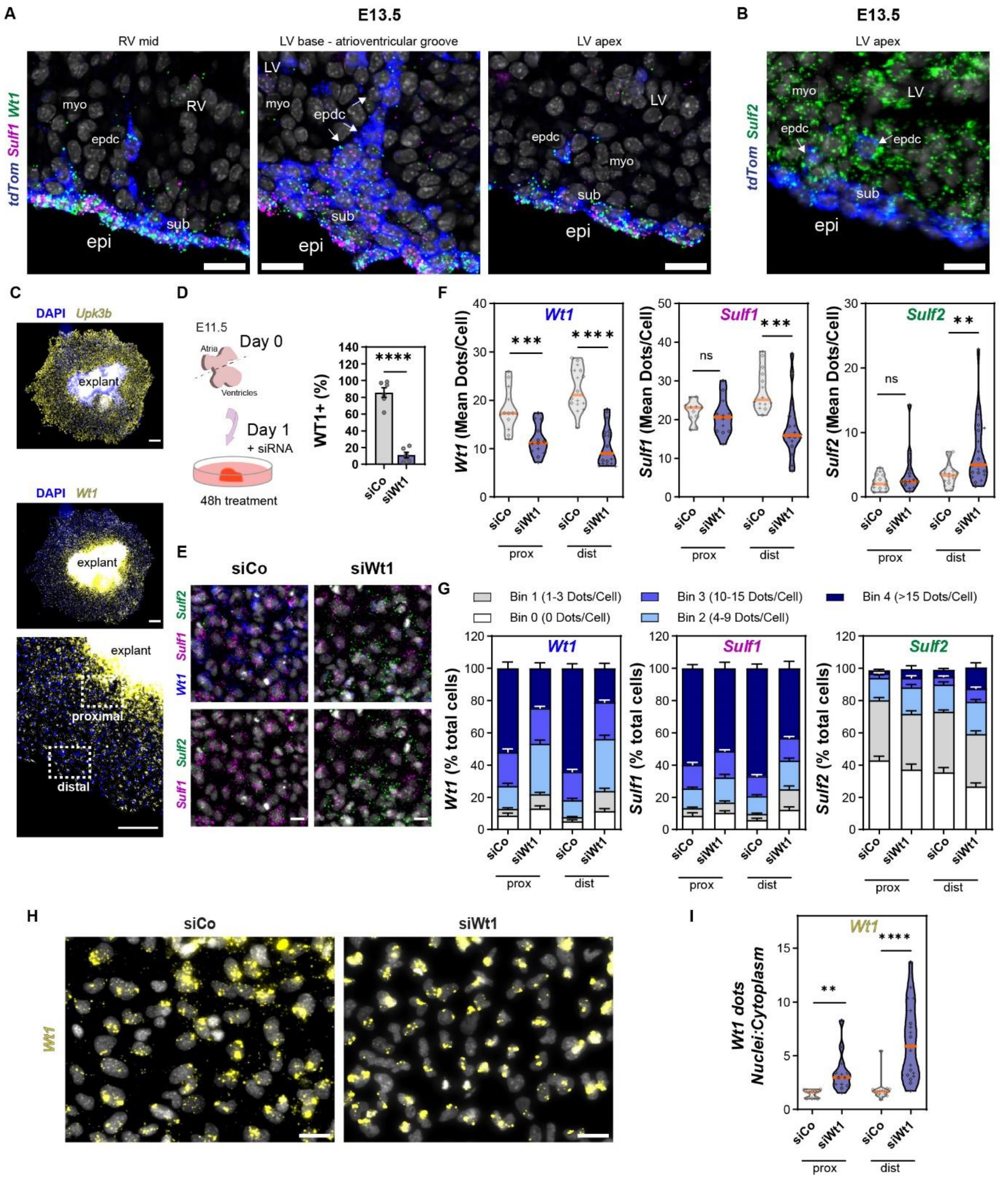

**Extended Data Fig. 5 Diminishing expression of *Sulf1* and *Wt1* coincides with epicardial EMT**

(A) Fluorescence ISH of E13.5 heart cryosections for *tdTomato*, *Sulf1* and *Wt1*, showing downregulated expression of *Wt1* and *Sulf1* in epicardial-lineage subepicardial (sub) and mesenchymal cells invading the myocardium (epdc). Scale bars: 20 µm. Images representative of n = 4 hearts. (B) Fluorescence ISH of E13.5 heart cryosections for *tdTomato* and *Sulf2*, showing upregulated expression of *Sulf2* in epicardial-lineage subepicardial (sub) and mesenchymal cells invading the myocardium (epdc). Scale bars: 20 µm. Images representative of n = 5 hearts. (C) Fluorescence ISH of E11.5 epicardial explants for *Upk3b* and *Wt1*, showing highest *Wt1* expression proximal to the explant piece. Scale bars, 200 µm.

(D) E11.5 epicardial explants transfected with control (siCo) or Wt1 siRNA (siWt1) for 48h. Quantification of percentage WT1 positive epicardial cells in control and Wt1 siRNA-transfected explants showing successful knock-down. Data are represented as mean ± SEM (n = 6 explants). Unpaired t-test (****) P <0.0001. (E) Fluorescence ISH of control and Wt1 siRNA-transfected epicardial explants for *Wt1*, *Sulf1* and *Sulf2,* and (F) corresponding quantification of expression levels. Violin plot symbol cross (+) represents atrial explant; diamond (◊) represents ventricular explant. Orange line indicates the median (n = 14-22 explants). Unpaired t-test with Welch's correction (**) P <0.01, (***) P <0.001, (****) P <0.0001 and ns, not significant. (G) Percentage of epicardial cells expressing *Sulf1* and *Sulf2*, grouped into 5 bins based on the number of dots (individual RNA molecules) per cell. Scale bars, 20 µm. Images representative of n = 13-19 explants. Data are represented as mean ± SEM (n = 13-19 explants). (H) Fluorescence ISH of control and Wt1 siRNA-transfected epicardial explants showing *Wt1* expression, with corresponding quantification of (I) *Wt1* transcript localisation across cell compartments. Scale bars, 20 µm. Images representative of n = 13-19 explants. Violin plot symbol cross (+) represents atrial explant; diamond (◊) represents ventricular explant. Orange line indicates the median (n = 13-19 explants). Unpaired t-test with Welch's correction (**) P <0.01, (****) P <0.0001.

*epi, epicardium; myo, myocardium; sub, subepicardium; epdc, epicardium-derived cells; RV, right ventricle; LV, left ventricle; prox, proximal; dist, distal.*

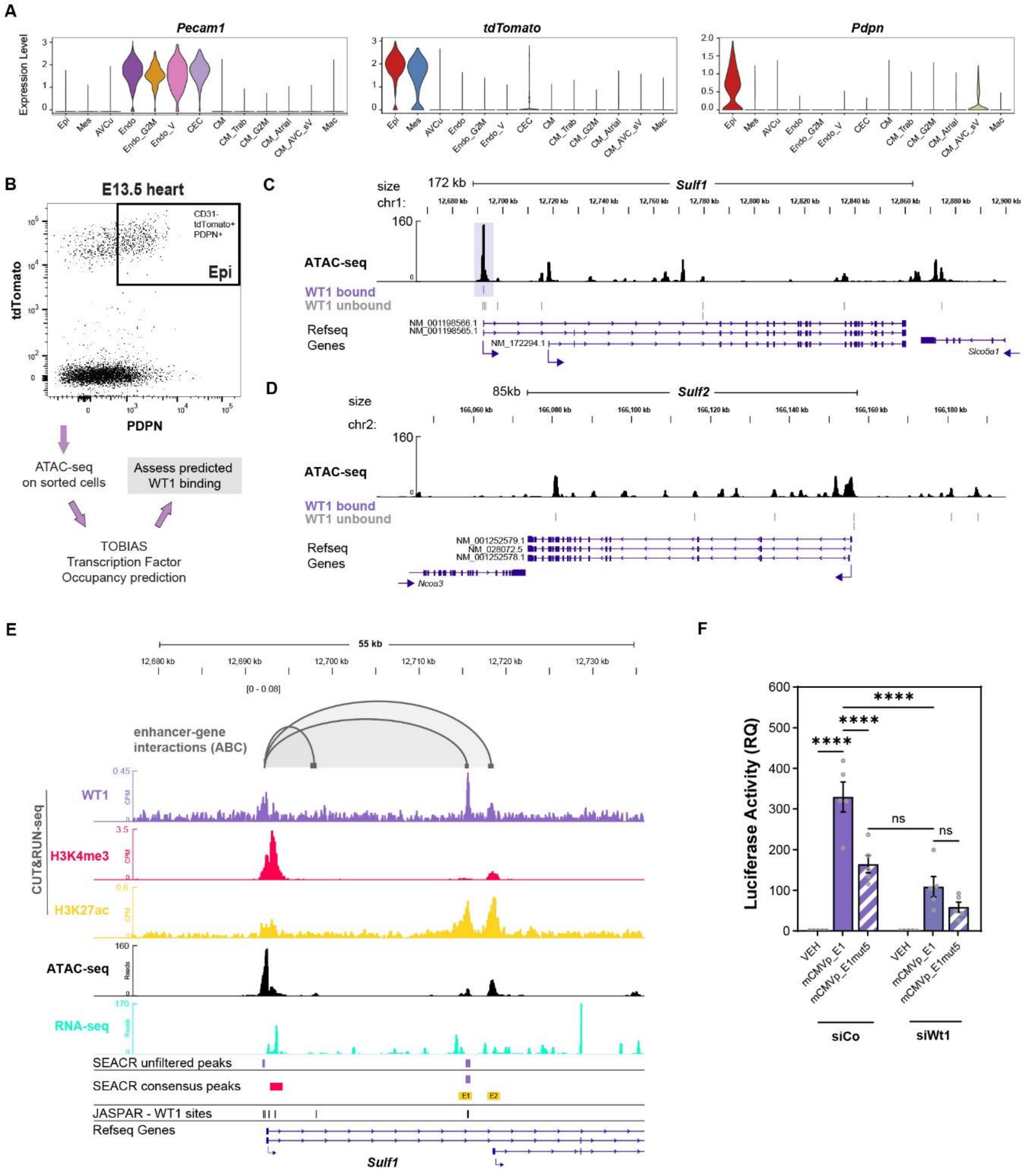


**Extended Data Fig. 6 Identification of WT1 associated CREs regulating *Sulf1* and *Sulf2* transcription in the epicardium**

(A) Violin plots showing relative expression of markers used to distinguish epicardium in E13.5 hearts. (B) Strategy used to gate and purify epicardial cells (CD31$^-$tdTomato$^+$PDPN$^+$) from E13.5 hearts. Purified epicardial cells were processed for ATAC-seq analysis, and downstream transcription factor occupancy prediction to assess binding of WT1 in the *Sulf1* and *Sulf2* loci. (C-D) Genome tracks showing predicted binding of WT1 at the (B) *Sulf1* and (C) *Sulf2* locus in sorted E13.5 epicardial cells. Track signal is in reads per million (scaled) and merged from n = 2 independent replicates. TOBIAS predicted WT1 binding calculated from n = 3 independent replicates (n = 2, E13.5; n = 1, E12.5). Region of interest (ROI) with bound WT1 indicated with purple shading. (E) Genome tracks of CUT&RUN-seq, ATAC-seq and RNA-seq showing direct binding of WT1 to intronic enhancer (E1) at the *Sulf1* locus of epicardial cells. Chromatin interaction tracks show linkages between WT1-bound enhancer E1 and upstream *Sulf1* promoter (ABC). All linkages shown have an ABC score >0.04. SEACR consensus peaks calculated from n = 3 (WT1, IgG and H3K27ac) and n = 4 (H3K4me3) independent replicates. JASPAR database WT1 binding sites indicated. (F) Epicardial cell line transfected with control (siCo) or Wt1 siRNA (siWt1) for 24h, followed by transfection with mCMVp_E1 and mCMVp_E1mut5 for 24h. Corresponding quantification of Luciferase activity to assess transcriptional activity of WT1 on intronic *Sulf1* enhancer in the epicardial cell line. Values expressed relative to vehicle control. Data are represented as mean ± SEM (n = 5 experiments). One-Way ANOVA and Šídák multiple comparison test (****) P <0.0001 and ns, not significant.

*AVCu, atrioventricular cushion; CEC, coronary endothelial cells; CM, ventricular cardiomyocytes; CM_Atrial, atrial cardiomyocytes; CM_AVC_sV, atrioventricular canal and sinus venosus cardiomyocytes; CM_Trab, trabecular cardiomyocytes; Endo, endocardial cells; Endo_V, valvular endothelial and endocardial cells; Epi, epicardium; G2M, proliferating; Mac, macrophages; Mes, mesenchyme.*

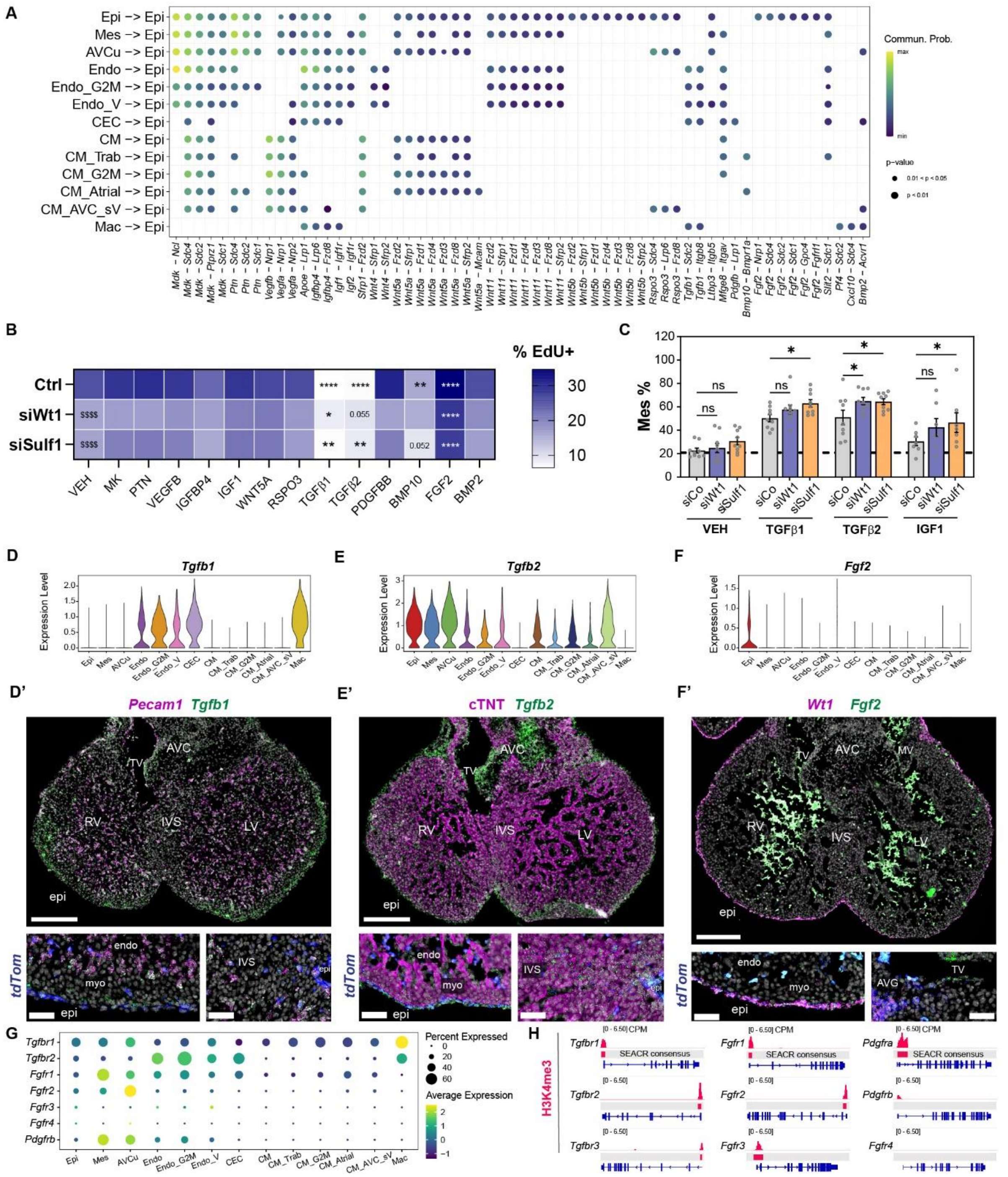

**Extended Data Fig. 7 Ligand-Receptor inference analysis identified HSPG-dependent incoming signalling and potential cell type sources in the E13.5 heart**

(A) Dot plot indicating source cell types communicating with the epicardium and corresponding key ligand-receptor pairs. High communication probability is indicated in yellow, dot size indicates p-values. (B) Epicardial cells transfected with control (siCo), Wt1 (siWt1) or Sulf1 siRNA (siSulf1) for 48h, followed by treatment with candidate cytokines for 24h, and corresponding quantification of percentage EdU positive cells. Data are represented as mean (n = 4 - 11 experiments). Two-Way ANOVA and Dunnett multiple comparison test (*) P <0.05, (**) P <0.01, (****) P <0.0001, and ($$$$) P <0.0001 (comparison between siRNA treatments + VEH). (C) Epicardial cells transfected with siRNAs for 48h, followed by treatment with candidate cytokines for 24h, and corresponding quantification of percentage mesenchymal cells. Data are represented as mean ± SEM (n = 7 – 9 experiments). One-Way ANOVA and Fisher's LSD test (*) P <0.05 and ns, not significant. (D) Violin plot showing relative expression of *Tgfb1* in hearts at E13.5, and corresponding (D') fluorescence ISH of heart cryosections for *Pecam1* and *Tgfb1*, showing majority source of *Tgfb1* from endocardial and coronary endothelial cells. Insets highlight proximity of epicardial-lineage cells (*tdTomato* positive) to *Tgfb1*-expressing endothelial cells. Scale bars: 200 µm and inset 50 µm. Images representative of n = 3 hearts. (E) Violin plot showing relative expression of *Tgfb2* in hearts at E13.5, and corresponding (E') fluorescence ISH of heart cryosections for cTNT and *Tgfb2*, showing widespread expression of *Tgfb2* throughout the ventricles. Insets highlight enriched expression in the epicardial layer and apical region of the intraventricular septum. Scale bars: 200 µm and inset 50 µm. Images representative of n = 3 hearts. (F) Violin plot showing relative expression of *Fgf2* in hearts at E13.5, and corresponding (F') fluorescence ISH of heart cryosections for *Wt1* and *Fgf2*, showing two specific sources of *Fgf2*, the epicardium and atrioventricular valves. Insets highlight localised expression in epicardial and valve endothelial cells. Scale bars: 200 µm and inset 50 µm. Images representative of n = 3 hearts. (G) Dot plot showing relative expression of growth factor receptors in hearts at E13.5. High expression is indicated in yellow, dot size indicates percentage expressed. (H) Genome tracks showing H3K4me3 profiles at *Tgfbr1, Tgfbr2, Tgfbr3, Fgfr1, Fgfr2, Fgfr3, Fgfr4, Pdgfra* and *Pdgfrb* locus. Track signal is in counts per million (CPM) and merged from n = 2 independent replicates. SEACR consensus peaks calculated from n = 4 (H3K4me3 and IgG) independent replicates.

*AVCu, atrioventricular cushion; CEC, coronary endothelial cells; CM, ventricular cardiomyocytes; CM_Atrial, atrial cardiomyocytes; CM_AVC_sV, atrioventricular canal and sinus venosus cardiomyocytes; CM_Trab, trabecular cardiomyocytes; Endo, endocardial cells; Endo_V, valvular endothelial and endocardial cells; Epi, epicardium; G2M, proliferating; Mac, macrophages; Mes, mesenchyme; epi, epicardium; myo, myocardium; endo, endocardium; RV, right ventricle; IVS, interventricular septum; LV, left ventricle; AVC, atrioventricular canal; AVG, atrioventricular groove; TV, tricuspid valves; MV, mitral valve.*
